# Lectins-glycoconjugates interactions: Experimental and computational docking studies of the binding and agglutination of eight different lectins in a comparative manner

**DOI:** 10.1101/2020.05.01.070102

**Authors:** Amit Kumar, Vijaya Kumar Hinge, Ashapogu Venugopal, Siva Kumar Nadimpalli, Chebrolu Pulla Rao

**Affiliations:** Bioinorganic Laboratory, Department of Chemistry, Indian Institute of Technology Bombay, Powai, Mumbai – 400 076, India; Department of Biochemistry, School of Life Sciences, University of Hyderabad, Hyderabad – 500 046, India

**Keywords:** lectin, glycoconjugate, hemagglutination, fluorescence spectroscopy, CD spectroscopy, molecular docking

## Abstract

Altering the lectin properties by chemically synthesized glycoconjugates is important in glycobiology. A series of eight plant lectins with varying carbohydrate specificity were chosen as model systems to study the binding by synthetic glycoconjugates. One of our earlier paper ^1^ deals with the binding of glycoconjugates by jacalin. Further to this, we have now extended the studies to several other lectins having specificities towards glucose/mannose, galactose and lactose, and the results are reported in this paper on a comparative manner. The binding aspects were established by hemagglutination and fluorescence spectroscopy, and the conformational changes by CD spectroscopy. Out of the fourteen glycoconjugates used in the present study, a galactosyl-naphthyl derivative, **1c** turns out to be most effective towards galactose-specific lectin in agglutination inhibition, fluorescence quenching by inducing considerable conformational changes. Similarly, mannosyl-naphthyl derivative, **3c** turns out to be most effective in inhibiting the agglutination of Glc/Man specific lectins. Present study demonstrates differential recognition of conjugates towards lectins. The results also supported the existence of a correlation between the glycoconjugate and lectin specificity at the carbohydrate recognition domain (CRD). The glycoconjugate that inhibits the agglutination binds in the CRD via polar interactions as well as by nonpolar/hydrophobic interactions arising from the aromatic moiety of the conjugate, whereas, the non-inhibiting conjugates bind primarily *via* hydrophobic interactions. The specific and selective binding of the glycoconjugates by these lectins were proven by the docking studies. Thus, the present study has contributed immensely towards understanding the molecular interactions present between the lectins and small molecules that will eventually help better drug design where the presence of hydrophoibic moieties would play important role.

## INTRODUCTION

Carbohydrate-protein interactions are involved in cell adhesion and cell communication processes that govern basically the social behavior of cells. Inflammation, tumor progression and metastasis, cell development and embryogenesis, and infection are some of the important biological events in which carbohydrates play important role by exhibiting interactions with the corresponding protein receptors.^2–6^ Lectins have been particularly useful in the study of molecular recognition of cellular oligosaccharides, owing to their ability to detect any variations in the structure of carbohydrates. Lectins are useful in isolation and characterization of the glycoconjugates.^7^ The substrates used for lectin studies are either mono- or oligo-saccharides because of their selective binding in the carbohydrate recognition domain. The interaction by the CRD can be utilized in order to modify the properties of lectins by appropriately derivatizing the carbohydrates.^1, 8^ In order to do that, the glycoconjugates should be modified so as to acquire hydrophobic character since many of the lectins have aromatic amino acids, such as, Trp, Tyr and Phe in their primary sequence and some of these are present in CRD.^1, 9–10^ Attachment of the aromatic group at C1 or C2 position of the glycoconjugate provides new direction of activity in modifying the properties of lectins as demonstrated by us recently in case of jacalin^1^, glycosidase^8^ and chemical nuclease^11^ and others^12^. Therefore, this paper deals with an extensive study of the interaction of aromatic-imino-glycoconjugates with eight different lectins possessing different carbohydrate specificities, *viz.,* galactose, glucose/mannose and lactose. These studies encompass both the experimental and computational methods and provide several rationalizations and the importance of hydrophobic interactions present between proteins and ligands that will have relevance in drug design and development.

## RESULTS AND DISCUSSION

The precursor glycose and its aromatic-imino-glycoconjugates have been used which were reported in our earlier studies ^1, 8^ and the schematic structures of these are given in Scheme 1. Eight lectins with different carbohydrate specificities, such as, galactose {pea nut agglutinin (PNA)^13^, soy bean agglutinin (SBA)^14^, *Dolichos lab lab* lectin II (DLL-II)^15–16^, *Moringa oleifera* lectin (MOL) ^17^}, glucose/mannose {pea lectin (PL) ^18^, lentil lectin (LL) ^19^, *Dolichos lab lab* I (DLL-I) ^15–16^}, and lactose {Unio lectin (UL)} have been isolated and purified using the affinity followed by gel filtration chromatography. Interactions of the glycoconjugates with purified lectins and possible inhibition of agglutination have been assayed using standard protocols ^1^. The binding and conformational changes have been addressed by fluorescence and CD spectroscopy. The interactions were modeled by the computational docking studies.

**SCHEME 1.**
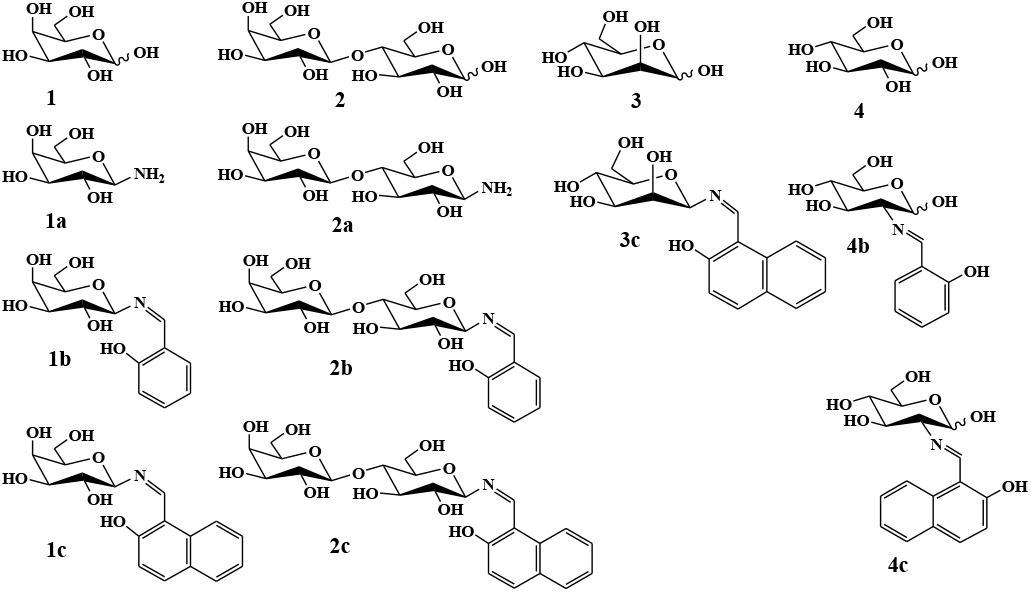
Line representation of glycoconjugates. ‘1’, ‘2’, ‘3’ and ‘4’ refers to simple galactose, lactose, mannose and glucose respectively. The conjugates ‘1a’, ‘1b’ and ‘1c’ are derived from galactose; ‘2a’, ‘2b’ and ‘2c’ are derived from lactose; ‘4b’ and ‘4c’ are derived from glucose and ‘3c’ is derived from mannose ^1, 8^.

### Hemagglutination assay

Lectins have been studied for their hemagglutination in the presence and in the absence of glycoconjugates by serial dilution method. Minimal agglutination inhibitory concentrations (MAIC) have been determined for all the cases and are listed in supporting information as Table S1. Relative inhibitory potency of the agglutination has been calculated by considering the potency of the simple precursor carbohydrate taken as one. It is dependent upon the lectin specificity for the carbohydrate, *e.g.*, DLL-II is a Gal specific lectin and the potency of Gal has been considered as one. The plots indicating the relative potency are given in Figure 1. It is evidenced from the plots that the glycoconjugates exhibit 3 – 100 times greater inhibition as compared to the corresponding simple glycose. The naphthylidine conjugates are better inhibitors than the corresponding amine or salicylidine conjugate.

**FIGURE 1.**
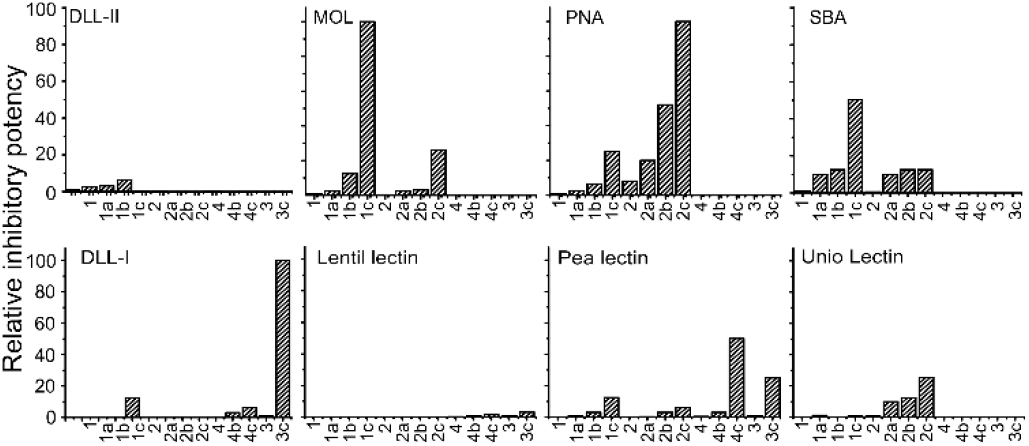
Relative inhibitory potency of different lectins by various glycoconjugates, where the inhibition by the corresponding glycose was taken as one.

The naphthylidine conjugate **1c** exhibited highest relative inhibitory potency in case of MOL, while SBA shows one half of this, and the PNA shows half of that of SBA and thus the same follows an order, MOL ≫ SBA ≫ PNA. In case of **2c**, the inhibitory potency is maximum with PNA, while the UL and MOL shows only one fourth of that of PNA. The **3c** is selective and shows maximum potency with DLL – I, while PL shows one-fifth of it. The **4c** shows a 50 folds relative inhibitory potency in case of PL. All this clearly shows that the relative inhibitory potency depends on the nature of the lectin possessing CRD and also the conjugate. Therefore, the data has been analyzed based on the nature of the lectin and that of the glycoconjugate as reported in this paper.

#### Comparison of the inhibition of agglutination based on the nature of lectins

Inhibition of agglutination has been analyzed for all the lectins based on their carbohydrate specificity. The Gal specific lectins (*viz.*, DLL–II, PNA, SBA, and MOL) are inhibited by the conjugates of Gal, particularly that of **1c**. This indicates that the carbohydrate moiety present in the glycoconjugate is specifically recognized by the CRD. Among the galactose specific lectins, MOL showed the lowest inhibitory concentration by **1c**, i.e., at 0.039 mM. Indeed the galactose specific proteins, viz., PNA, SBA, and MOL, were also inhibited by the lactose counterpart, i.e., **2c**, but at a greater concentration (i.e., 0.15±0.01 mM) than that exhibited by **1c**. This indicates that these lectins also recognize the galactose moiety of the lactose. All these results reflect on the difference between the conjugate of monosaccharide and that of a disaccharide. Similarly, the lactose conjugate (**2c**) inhibits the lactose specific lectin, i.e., UL at much lower concentration, such as, 0.019 mM and to some extent by the conjugate of galactose (**1c**), i.e., at 0.62 mM. On the other hand, glucose or mannose derivatives do not inhibit the UL even upto 10 mM. Activity of all glucose/mannose specific lectins was inhibited only by the conjugates of glucose and mannose and not by those of galactose or lactose (Figure 1). Among the glucose/mannose lectins, DLL-I and LL showed specific interaction towards the recognition of glucosyl- and mannosyl-conjugates. The activity of LL is inhibited by the glucosyl-conjugates at an equal or higher concentration than that of the conjugates of mannose, viz., 10±0.5 (**3a**) and 20±0.5 (**3b**) *vs.* 5±0.01 (**4c**) mM. However, the PL showed non-specific behaviour and its activity was inhibited by all the glycoconjugates. Notably, the MAICs were lower in case of glucosyl- and mannosyl-conjugates. The results suggested the specific recognition of the glycoconjugates at the CRD, besides exhibiting other interactions. Among the eight lectins studied, no considerable inhibition of agglutination was observed in case of DLL – II, LL and UL. However, greater inhibition of agglutination was observed in case of PNA, MOL and DLL – I, while SBA and PL showed intermediate nature.

#### Comparison of inhibition of agglutination based on the nature of glycoconjugates

The inhibition of the agglutination activity of each of the lectin has also been analyzed based on the nature of the glycoconjugates. Galactose (**1**) and its conjugates mainly inhibit the Gal/Lac specific lectins. The MAIC’s follows a trend, *viz.*, **1c** > **1b** > **1a** > **1**, for the Gal specific lectins and it varies from 3.9 to 125, 0.39 to 25, 0.31 to 5 and 0.04 to 1.25 mM respectively for **1**, **1a**, **1b** and **1c**. Similar behavior was exhibited even by other conjugates, where the inhibition is dependent upon the lectin specificity. For example, Lac and its conjugates inhibit the agglutination mainly of the Lac and Gal specific lectins and not the Glc/Man specific ones. Among all the lectins, PL showed nonspecific interaction and its activity was inhibited by all the conjugates at concentrations ranging from 0.31 to 500 mM, but the effect on inhibition is rather minimal. It is interesting to note that even though the PL showed the nonspecific interaction, its activity was mainly inhibited by the conjugates possessing Man and Glc units. The presence of hydrophobic moiety, *viz.*, naphthyl or salicilyl, makes them effective inhibitors so as to exhibit 100 – 200 times better inhibitory potency than the corresponding simple carbohydrate. For all these lectins, the naphthylidene-glycoconjugate inhibits agglutination at a lower concentration than the corresponding salicylidene ones. Such conjugates inhibited better than the C1-amine as well as simple glycoses. Thus, it is interesting to note that the naphthyl-conjugates always showed higher inhibition than the corresponding salicyl- or simple amine based conjugates. In general, the inhibitory potency follows a trend, viz., simple carbohydrates < C1/C2-amines of glycoses < salicyl-imino-conjugates < naphthyl-imino-conjugates. The trend thus observed among the glycoconjugates was similar in case of all the lectins.

### Fluorescence spectroscopy

Fluorescence spectroscopy is an ideal technique to analyze protein – small molecular interactions. Fluorescence spectra showed protein intrinsic emission at λ_max_ 335 – 340 nm depending upon the lectin. The observed spectra exhibits a decrease in the fluorescence intensity (fluorescence quenching) and this is a function of the concentration of the glycoconjugate.

#### Fluorescence titration of lectins with glycoconjugates

Representative spectra are shown for SBA and PL in Figure 2 on going from simple glycose to amine to salicylidene to naphthylidene derivate. Each of the LL and PL possesses ten Trp residues, of which eight are well buried inside the protein and the remaining two are present on the surface. Of the 12 Trp residues present in PNA, four are exposed and the remaining are buried inside the protein. Of the 24 Trp residues present in SBA, 16 are buried and the rest eight are exposed. Careful examination of the fluorescence spectra shows the presence of two different classes of Trp (buried *vs* exposed) (Figure S1 – S8) and is in agreement with that observed from the crystal data (e.g., LL ^20^, PL ^21^, PNA ^22^ and SBA ^23^). Corresponding relative quenching curves were shown for each protein in the supporting information under Figure S10.

**FIGURE 2.**
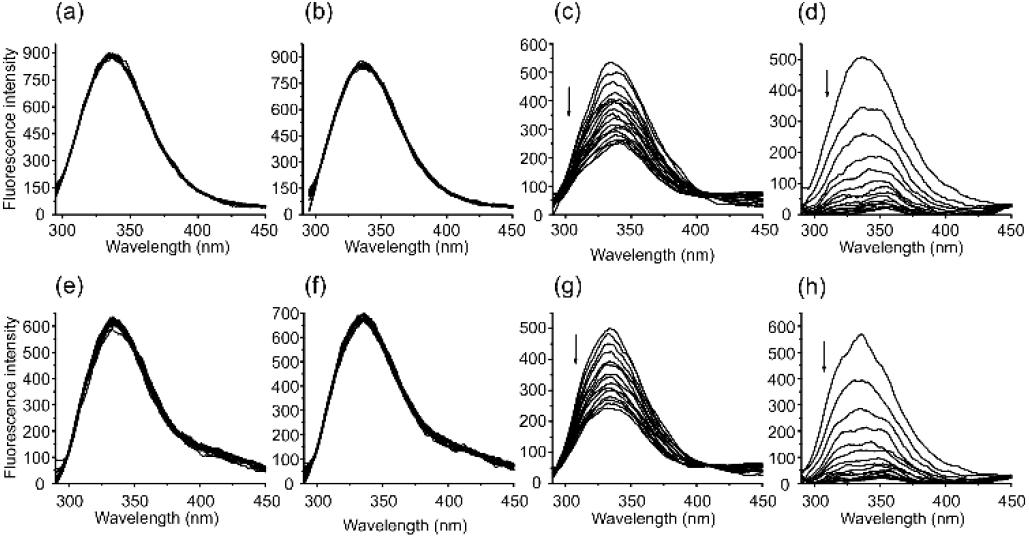
Fluorescence spectral traces obtained during the titration of lectins by glycoconjugates: SBA by (a) **1**, (b) **1a**, (c) **1b** (d) **1c**; and pea lectin (PL) by (e) **1**, (f) **1a**, (g) **1b**, and (h) **1c**. Arrow direction indicates fluorescence quenching.

Intensity of both the Trp was diminished upon the titration with naphthylidene glycoconjugates, of which λ_em_^max^ of one Trp seems to remain at 335 nm while the other shows a red shift towards 350 nm (Figure 2d and 2h), suggesting that the environment of one type of Trp was changed in presence of the glycoconjugate. The red shift indicates that the lectins undergo conformational changes upon interaction with glycoconjugates that brings the Trp residues onto the surface of the protein.

Maximum quenching in fluorescence intensity was observed only in case of naphthyl derivatives. This suggested, that immaterial of the carbohydrate specificity of the lectin, **1c** always exhibits high fluorescence quenching, as compared to the corresponding salicylidene derivative. However, fluorescence quenching was not observed in case of simple carbohydrates (**1**, **2**, **3**, **4**) or their C1 amines (**1a**, **2a**) (Figure 2 & Figure S1 to Figure S9). This suggested that even if the glyco-part is not recognized by the corresponding lectin as can be noticed from the agglutination data given in Figure 1 and Table S1, the conjugate interacts with the lectin using its aromatic moiety. It is interesting to note that other conjugates (**2c**, **2b**, **4c**, **4b**) were able to quench the fluorescence intensity, but they are not able to inhibit hemagglutination property, as the glyco-portion differs from that of the lectin specificity. For example, **2c** quenches the intensity of glucose/mannose specific lectins, *viz.*, DLL-I, DLL-II and LL, but unable to inhibit the agglutination. This indicates that carbohydrate is being recognized and bound at the CRD; and the aromatic moiety plays important role in binding with the protein.

**TABLE 1.** Calculated IF_50_ values based upon the relative intensity plots which were obtained when glycoconjugates were titrated with the lectins. Table foot notes same as in Table 1. In case of the compounds, 1, 1a, 2, 2a, 3 and 4 the IF_50_ could not be determined.

| Conjugate | $IF_{50}$ (μM) | | | | | | | |
| --- | --- | --- | --- | --- | --- | --- | --- | --- |
|  | DLL-I | LL | PL | MOL | SBA | DLL-II | PNA | Unio |
| **1c** | 47±2 | 52±2 | 50±3 | 49±1 | 45±2 | 56±5 | 47±2 | 72±6 |
| **1b** | 207±8 | 180±5 | 333±23 | 152±7 | 335±33 | 260±24 | 307±18 | 330±24 |
| **2c** | 86±4 | 250±7 | 480±19 | 59±1 | 414±32 | 283±19 | 366±16 | 430±21 |
| **2b** | 87±3 | 122±6 | 100±8 | 124±12 | 112±2 | 164±8 | 144±11 | 105±8 |
| **3c** | 132±9 | 91±2 | 171±5 | 117±4 | 166±13 | 122±12 | 123±7 | 91±3 |
| **4c** | 250±9 | 133±10 | 305±20 | 278±11 | 279±22 | 167±7 | 181±10 | 276±25 |
| **4b** | 317±5 | 345±12 | 371±11 | 350±21 | 384±32 | 370±35 | 396±24 | 363±29 |

#### Extent of fluorescence quenching

The extent of fluorescence quenching has been further gauzed by comparing the concentrations at which 50% of the fluorescence is being quenched and this is defined as IF_50_. The IF_50_ (μM) values were derived from the relative intensity plots and the corresponding data is given in Table 1 (Figure S9). The observed IF_50_ values are in the range of 50 to 300 μM for naphthylidene-conjugates and 90 to 500 μM for salicylidene-ones showing a two folds higher effect on protein by the naphthylidene conjugates. This difference is at least four-fold when **1c** is compared with **1b**. This is attributable for the difference between phenyl and naphthyl moieties present in these and their effective interaction with the protein. Further, the trends in the quenching behavior have been analyzed from the point of view of the lectins as well as the glycoconjugates and representative ones are given in Figure 3 and the plots for all the cases are given in the supporting information under Table S2, S3 and Figure S10. For full instructions, please see the journal’s Instructions for Authors.

**FIGURE 3.**
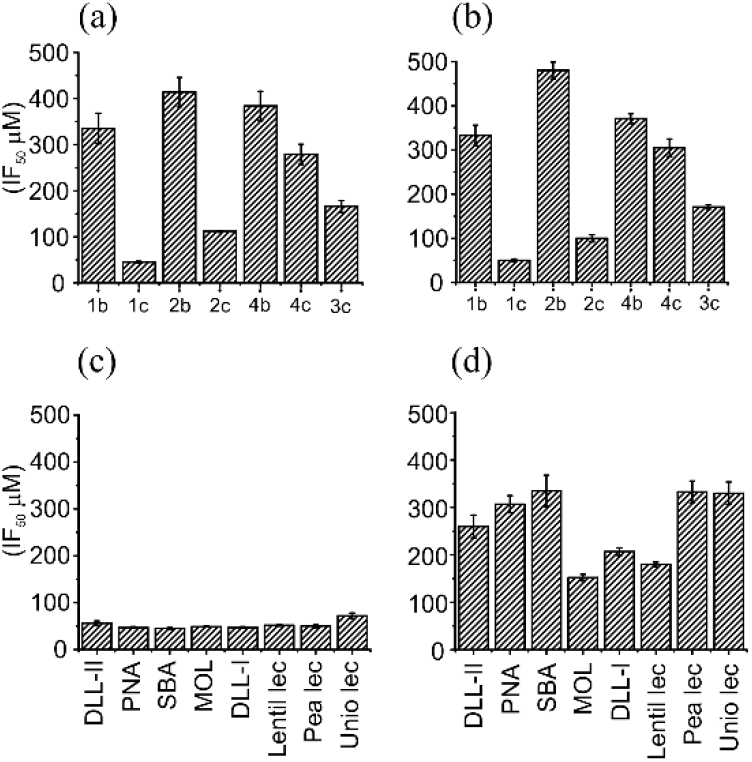
IF_50_ (μM) plots for SBA *vs.* glycoconjugates in (a) and PL *vs.* glycoconjugates in (b). Histogram of IF_50_ (μM) for **1c** *vs.* lectins is given in (c) and that for **1b** *vs.* lectins in (d).

Considering lectins, it was observed that naphthylidene conjugates always showed low IF_50_ values as compared to the salicylidene ones. The general behavior of fluorescence quenching is independent of the glycose. However, there are significant differences in IF_50_ even among the salicylidene or naphthylidene conjugates. Thus, it highlights the importance of glycoses and their recognition at CRD. Among the lectins, the glycoconjugates follow the general trend of, naphthylidene conjugates > salicylidene conjugates >>>glycose-NH_2_ ≥ glycoses. Considering the conjugates, the naphthylidene ones showed almost identical binding towards different lectins, however the salicylidene ones showed a broad range of IF_50_ values. Representative trends for galactose based conjugates are as follows, **1c**: DLL-I = MOL = SBA = PNA > LL ~ PL ~ DLL-II > UL and for **1b**: MOL > LL > DLL-I > DLL-II > PNA > PL ~ SBA ~ UL. The naphthylidene conjugates bind 2 – 4 times more efficiently as compared to the corresponding salicylidene conjugates (Figure 3). The trends observed in the fluorescence data clearly reveal that the glyco-imino-aromatic-conjugates bind lectins *via* carbohydrate and/or hydrophobic moiety at the CRD or the hydrophobic region or both. Although there are significant differences among conjugates for a particular lectin, in general the naphthylidene conjugates quenche the fluorescence better than the others in case of all the lectins. In case of few lectins, IF_50_ is same implying that the hydrophobic moiety plays dominant role over the glyco-moieties, and hence the glyco- specificity is being overruled in such cases.

#### Stern-Volmer analysis

Fluorescence quenching data for all the compounds used in this study were analyzed by Stern-Volmer equations (given in experimental section). The data corresponding to PL and SBA are given in Figure 4. All the data pertinent to the Stern-Volmer plots are given in the supporting information as Figure S11 and Table S4. In case of all the lectins, it is clearly seen that the naphthylidene conjugates have higher Stern-Volmer constant (*K*_*sv*_) than the corresponding salicilyl derivatives. A factor of 7–10 increase in *K*_*sv*_ was observed among the conjugates of **1** on going from salicilyl to naphthyl. However, this is only a factor of two differences in case of the conjugates of **2** and **4** towards the corresponding lectins. Similar trend was observed in case of *f*_*a*_ valve, where the naphthyl conjugates access the fluorophore more than the salicilyl derivatives. This data clearly suggests the previous observation that the naphthyl derivatives bind better than the salicilyl ones and hence be able to quench the fluorescence completely whereas the salicilyl derivatives are not. Fluorescence quenching follows a trend similar to previous observation, *viz.*, glycoses < glyco-C1-NH2 ≪< glyco-salicyl-conjugates < glyco-naphthyl-conjugates.

**FIGURE 4.**
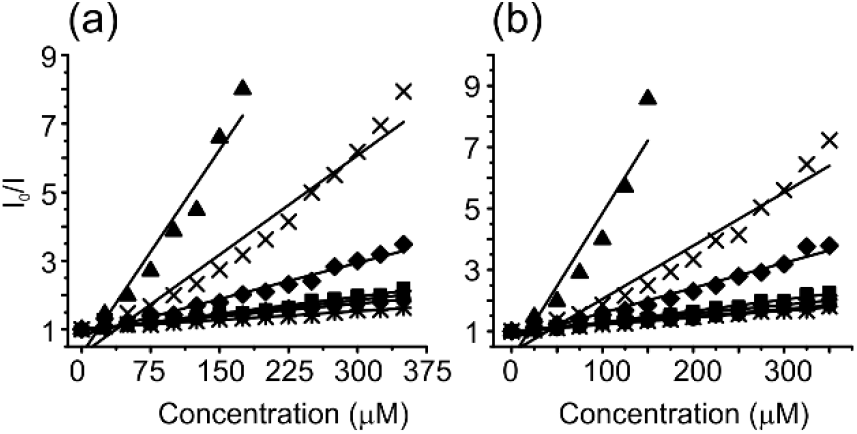
Stern-Volmer plots of pea lectin (PL) in (a) and SBA in (b) when titrated with various glycoconjugates. The symbols are same for both the plots: ★ – **1b**, ▲ – **1c**, × – **2b**, ✳ – **2c**, ◆ – **3c**, ⚫ – **4b** and ■ – **4c**.

The fluorescence results suggest that the nature of the carbohydrate and the derivatization at C1- or C2-are important in their differential interactions with lectins. Among the naphthyl derivatives, **1c** is a better quencher while it is **2b** in case salicilyl ones (Figure 4). The quenching behaviour of **2b** is similar to that of **4c**, indicating that the presence of disaccharide can be compensated by the addition of a higher hydrophobic moiety. The quenching of **1b** is similar to that of the **4b**. In effect, the naphthyl-conjugates are efficient quenchers when compared to their salicilyl counterparts and the C1-glycoconjugates were better quenchers than the C2-ones. This trend is same for different lectins as reported in this paper.

#### Circular dichroism (CD)

CD spectra were recorded in far UV region in the absence and in the presence of glycoconjugates in 5 mM Tris buffer, pH = 7.4. The spectra of native lectins, *viz.*, DLL-I, lentil, pea and PNA exhibit a minimum at 220 nm that is characteristic of β-sheet structure, whereas the other lectins, *viz.*, MOL and DLL-II exhibited characteristic band for α–helix. It has been reported that no conformational change or disruption in aromatic side chain of protein was observed in the presence of simple carbohydrates for many of the lectins or glycosidase ^1, 8–10^. Similarly, no change in ellepticity was observed when different lectins were titrated with glycoses or glycose-C1/C2–NH_2_, suggesting that these do not affect the secondary structure of the lectins. However, the introduction of aromatic moiety results in considerable changes in the ellipticity (Figure 5 and Figure S12 – S17).

**FIGURE 5.**
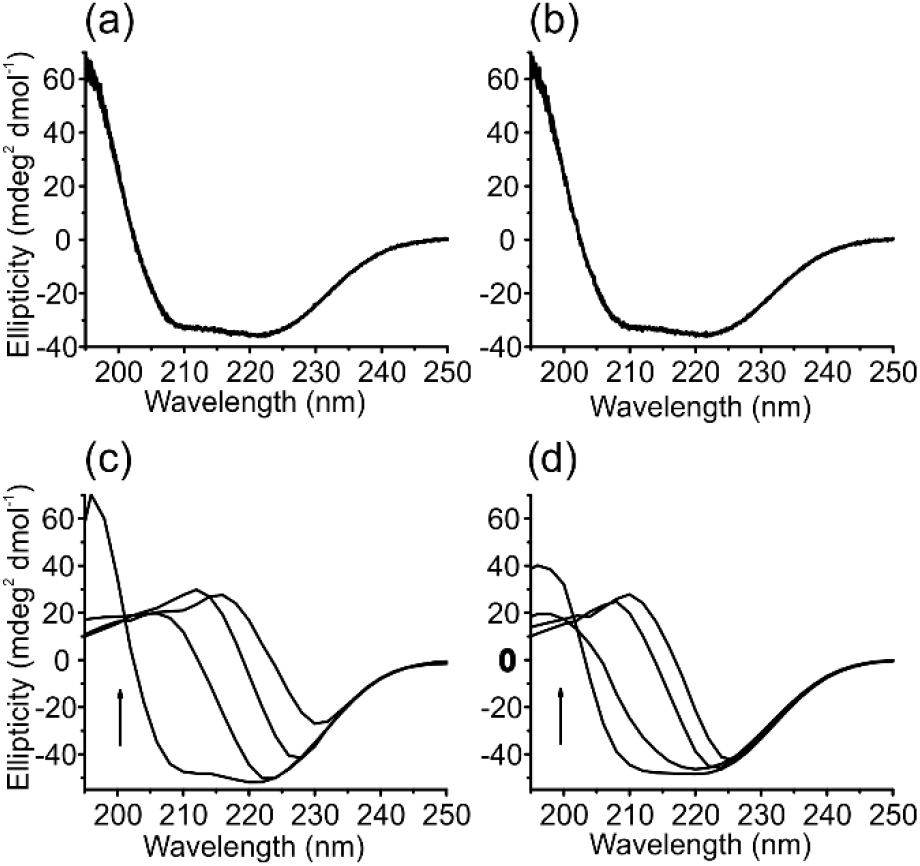
CD spectral traces obtained when MOL was titrated with **1** and its derivatives. (a) **1**, (b) **1a**, (c) **1b**, and (d) **1c**. Direction of arrow indicates conformational changes occurred in the protein.

The conclusions drawn from CD are in agreement with that drawn based on fluorescence studies. Large conformational changes and ~5 nm red shift in the band position were observed in the presence of glyco-imino-aromatic-conjugates. Naphthyl conjugates showed large change in the ellipticity as compared to the salicyl derivatives. The CD studies also suggest the requirement of hydrophilic as well as hydrophobic moieties simultaneously, in order to bring considerable changes in the ellipticity. It is **1c** that shows larger conformational changes as compared to all the other conjugates. This indicate that the glyco-moieties are important for the recognition at CRD, and higher the size of the aromatic moiety the greater the interactions and brings large conformational changes in proteins. Similar results were observed with the other conjugates of same glycose, *viz.*, **2c** *vs.* **2b**, **4c** *vs.* **4b** (Figure S12 to Figure S17).

All glyco-imino-aromatic-conjugates seem to perturb the secondary structure of lectins but unable to inhibit the agglutination. A few conjugates bind at the CRD and interacts with the aromatic side chain of amino acids resulting in ellipticity changes. However, the binding of **1**, **1a**, and **2a** do not bring such changes owing to the absence of the aromatic moiety in these, but these inhibit the agglutination by binding at the CRD. On the other hand, the glycoconjugates which were not inhibiting the agglutination are still able to change the ellipticity due to the presence of their aromatic moiety. These glycoconjugates, possibly, interact through hydrophobic patches present on the surface of the lectins and thereby cause conformational changes. However, the conformational changes brought about by these are not sufficient enough to disrupt the CRD to prevent lectins from their interaction with RBCs. So, agglutination occurs in the presence of these conjugates, even at a concentration that is 4 – 5 times higher than those which inhibit the agglutination. This means, that the aspects such as the nonspecific binding (π…π interaction), quenching observed in the fluorescence intensity, changes observed in the conformation of protein may have direct implications on the carbohydrate recognition or its agglutination property of the lectin.

### Docking studies

In order to identify the binding regions as well as the nature of interactions present between the glycoconjugates and lectins, docking studies were performed using Autodock software. All the derivatives were optimized using B3LYP/6-31G and were used for the docking. The corresponding docked regions show that the glycoconjugates occupy CRD (Figure 6, Figure S18 and S19).

**FIGURE 6.**
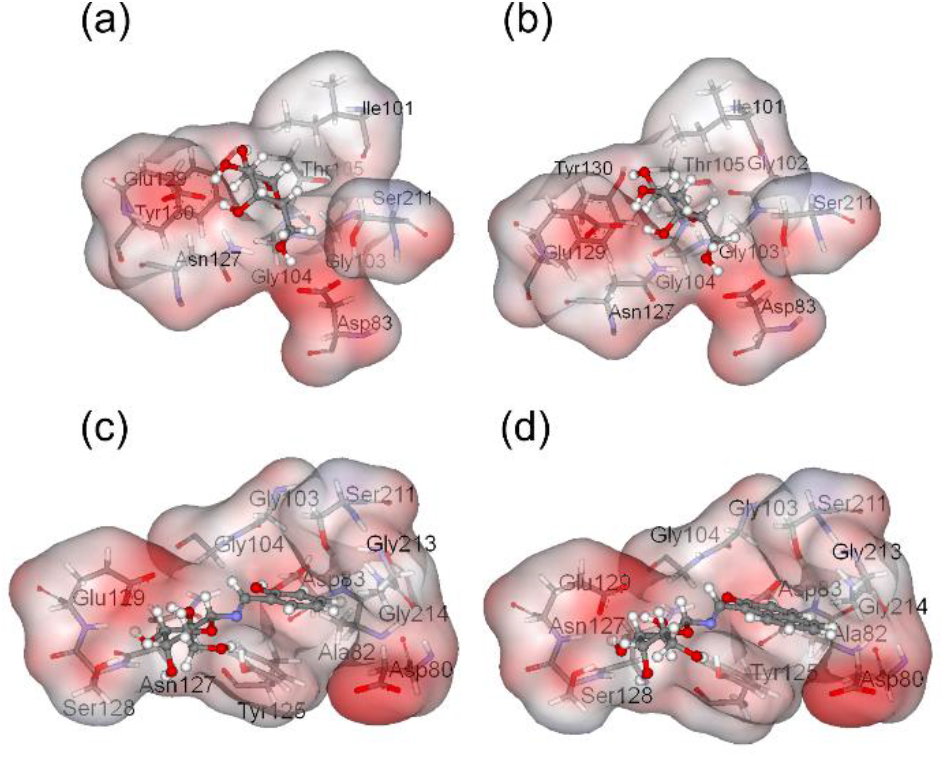
van der Waal surface representations (colored with electrostatic potential) showing the binding of glycoconjugates in case of PNA at CRD with (a) **1**, (b) **1a**, (c) **1b** and (d) **1c**.

In case of lentil lectin, the benzyl moiety of Phe123 shows π…π stacking with phenyl moiety of **4b** with a centroid to centroid distance of 4.2 Å. In case of PNA, π…π interaction was observed between **1b** and Tyr125 at a distance of 4.1 Å. The side chain of Ile101 of PNA shows C-H…π interaction with the phenyl moiety of **1b** at 3.6 Å. In SBA, the salicilyl imine moiety of **1b** and **2b** shows interaction with aromatic amino acids. The **1b** shows hydrophobic interaction with Tyr 107 and Ile 214 residues. The napthyl ring possessing glycoconjugates, **3c** and **4c** shows higher binding energy with lentil lectin due to the additional stabilization contributed from its aromatic moiety (Figure 6). This includes the π…π interaction contributed from the naphthyl ring of **3c** with benzyl side chain of Phe 123 at a centroid to centroid distance of 3.9 Å whereas in the **4c** the naphthyl ring is placed in the hydrophobic core formed between the Tyr 100, Trp 128 residues. This brings additional stabilization from the N-H…π interaction arising from Asn 125 and naphthyl moiety of **4c**. In PNA, the naphthyl ring of glycoconjugate **1c** is stabilized through π…π interaction with hydroxy phenyl side chain of Tyr 125 is at a centroid to centroid distance of 3.9 Å. The napthyl moiety of **2c** surrounded by the hydrophobic cluster formed by the Ile 101, Tyr 130 residues, and N-H…π interaction (3.9 Å) formed between Gly 104 and napthyl ring. In SBA, the naphthyl ring of **1c** shows interaction with the hydrophobic residues, Tyr 107, Trp 132, whereas the naphthyl ring of **2c** stabilized through π…π interaction with benzyl moiety of Phe 128 at distance of 4.2 Å.

At the docked regions, the interactions present between the aromatic moiety of glycoconjugates with the lectins were closely analyzed as given in Figure 7. Varying number of interactions exist between the conjugate and the lining amino acids of CRD depending upon the nature of the glyco-conjugate. For example, in case of lentil lectin, (Asp 81, Asn 125, Ala 30, Glu 31); (Ala80, Gly 99, Phe123); (Tyr 100, Gly 29); (Gly 98, Ala 127, Trp128); and (Gly 79, Ala 126) showed 5, 4, 3, 2, and 1 interactions with glycoconjugate respectively. For example, Ala80 of LL showed only one interaction with all conjugates while Glu31 of LL showed 7 interactions with **4**, **4b** and **4c** (Figure 7a). Similarly, in case of PNA, a maximum of 8 interactions were observed in case of **2**, **2b** and **2c** while only one interaction was seen in case of Ala 82 with **2**. A similar behavior was noticed even in case of SBA (Figure S18 and S19). Thus, different lectins showed varying number of interactions ranging from 15 to 32 depending upon the conjugate. Similarly, lentil lectin showed interactions with **3** and **4** (Figure 7b). This also supports the observation on lectin specificity towards the glycoconjugates as analyzed by hemagglutination. The analysis of interaction indicates that the salicilyl and naphthyl imines show better interaction through aromatic residues present on the lectins. The number of interactions as well as the contributions to the binding energies is higher for the aromatic portion of the glycoconjugate rather than the glycose-part of the same (Figure 7c). Thus the binding energies follow a trend, viz., simple sugar < sugaramines < salicilyl imines < naphthyl imines, suggesting that the imino-aromatic conjugates interact better at CRD of lectin owing to the presence of their imine and aromatic moieties as compared to the simple sugar or its C1-amine that is devoid of such moieties.

**FIGURE 7.**
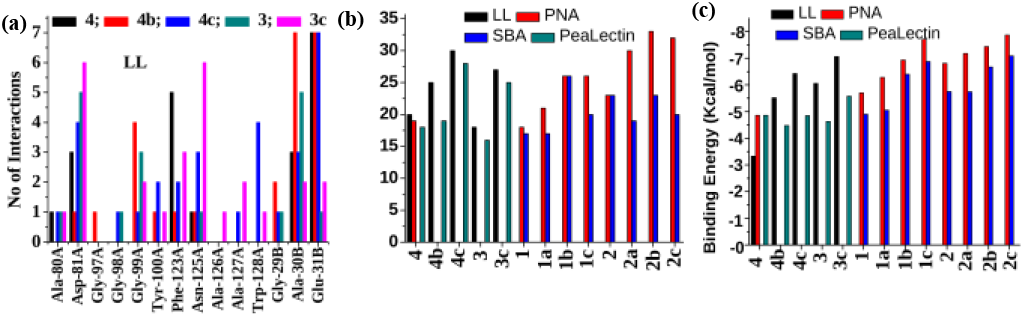
Analysis of the binding parameters, such as, number of interaction and binding energy, of the glycoconjugates when docked with the lectins: (a) total no of interactions exhibited by individual residues at CRD of LL towards particular glycoconjugate, (b) total number of interactions present between the glycoconjugate and the lectin residues at CRD based on the glycoconjugates, and (c) the calculated binding energies (kcal/mol) of the glycoconjugate by the corresponding lectin.

## CONCLUSIONS AND CORRELATIONS

Protein purified from the affinity chromatography followed by the gel filtration resulted in single band on the SDS-PAGE, indicating that the lectin is pure. Purified protein agglutinates RBCs giving further support for the presence of lectins and their purity. Agglutination inhibition exhibited by the glycoconjugates is in line with the lectin specificity. For example, Gal specific lectins *viz.*, DLL-II, PNA, SBA, and MOL are mainly inhibited by the conjugates of **1** and **2**. This indicated the selective recognition of the glycose moiety at the carbohydrate recognition domain. There were significant differences in the inhibitory potency between the glycoconjugates, particularly, ongoing form C1-amine derivatives to naphthyl ones. Pea lectin exhibit nonspecific behavior towards recognition of conjugates and thus its activity was inhibited by all the conjugates followed by a marginal difference in the inhibitory potency among the derivatives.

In general, it was observed that the naphthyl conjugates inhibited the lectin agglutination at lower concentration as compared to the other. Agglutination follows a trend among the glycoconjugates, viz., naphthyl derivatives > salicyl derivatives ≫ C1/C2 derivatives > simple carbohydrates. These results suggested a specific recognition of glycomoiety at the CRD and also importance exhibited the importance of the aromatic moiety.

The naphthyl modified glycoconjugates quenches the fluorescence intensity better when compared to salicilyl ones. However, no changes were observed in case of C1 derivatives or simple carbohydrates. Further, the naphthyl modified derivatives have higher Stern-Volmer constant as compared to the salicilyl derivatives, suggesting that the naphthyl-conjugates have higher affinity towards lectins. This is attributable to the larger size of the aromatic moiety present in these. Similar observations were noticed even in the agglutination studies where the naphthyl conjugates were found to be more effective. During the fluorescence titration, a red shift was observed in the λ_em_ that is attributable to protein conformational changes. Indeed it was observed that the lectins undergo conformational changes when titrated with glycoconjugates. These results are in agreement with agglutination and fluorescence studies, where naphthyl derivatives cause higher conformational changes than salicilyl ones, while C1/C2 derivatives or simple glycoses do not exhibit any conformational changes. The possible modes of binding of glycoconjugates have been given in Scheme 2 as a cartoon presentation in case of galactose binging lectin.

**SCHEME 2.**
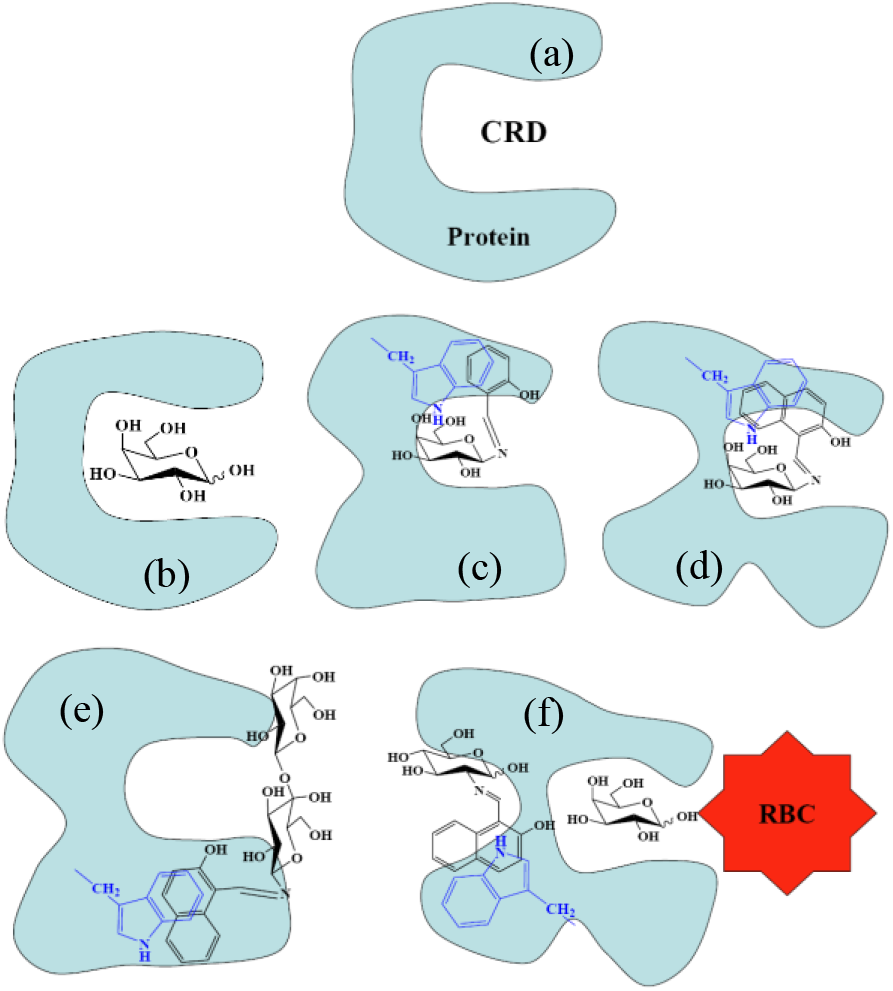
Cartoon representation of different modes of binding of glycoconjugates with a specific lectin. (a) Native protein exhibiting the CRD; (b) binding of simple glycose at the CRD that leads to the inhibition of agglutination and but brings no change in the protein conformation; (c) binding of salicyl derivatives which fit at the CRD and interacts with the proximal Trp and brings changes in the protein conformation; (d) naphthyl derivative causing greater conformational changes in the protein; (e) non-specific binding of disaccharide conjugate that induces conformational changes; (f) different modes of glycoconjugate binding that is away from CRD but causes conformational changes. In such cases, the CRD is intact and the agglutination occurs in presence of those conjugates at 5 – 10 times higher concentration.

The present study provided sufficient evidence for the differential binding of the naphthyl vs. salicilyl, C1-vs. C2-conjugation, specific recognition of the glycoconjugates at the CRD, and the contribution of hydrophilic vs. hydrophobic interactions with lectins. This further suggests that the changes observed in the lectin property by virtue of the binding of the glycoconjugate is not limited to one lectin but extended to several other lectins. All these add to the understanding the molecular interactions present between the lectins and small molecules. Therefore, the present study would greatly enhance the understanding of drug design as the introduction of the hydrophobic moieties (*viz.*, naphthyl or salicyl) into the glycoconjugate would indeed enhances their affinity towards lectins.

## EXPERIMENTAL SECTION

### Extraction and purification of lectins

Lectins were extracted from the seeds and the protein was purified by affinity followed by gel filtration chromatography galactose (pea nut agglutinin (PNA) ^13^, soy bean agglutinin (SBA) ^14^, *Dolichos lab lab* lectin II (DLL-II) ^15–16^, *Moringa oleifera* lectin (MOL) ^17^), glucose/mannose (pea lectin ^18^, lentil lectin ^19^, *Dolichos lab lab* I (DLL-I) ^15–16^. Brief of these are given here. All the lectins were extracted and purified by affinity chromatography followed by gel filtration chromatography. Precipitated fraction of the overnight extract was dialyzed and loaded on an appropriate affinity column (e.g., PNA on galactose affinity, LL on mannose specific column), pre-equilibrated with equilibration buffer (25 mM Tris, pH = 7.4). The outcome was reloaded on the column for 5 – 6 times successively. A fraction of 3 ml was collected, pooled and their purity was checked on SDS-PAGE. Gel filtration chromatography was performed on sephacryl S–100 column. Fractions of 1.5 ml were collected. These samples were analyzed on 12 % SDS-PAGE. The fractions, which gave the pure protein, were pooled and concentrated. The purified protein was used for further studies.

### Hemagglutination assay

Agglutination experiments were carried out in two steps according to the known procedure ^1^. In the first step, fresh rabbit RBCs were prepared by treatment of trypsin/pronase and in the second step, the agglutination was performed by serial dilution method. To the first well 50 μl (1mg/ml) of lectin was added and subsequently serially diluted. To study the inhibitory potency of the glycoconjugates, these were serially diluted and a fixed amount of lectin was added. This mixture was incubated for 15 min at 37 °C. Subsequently, fixed amounts (50 μl) of RBCs were added, and the observations were recorded after 90 minutes of incubation.

### Fluorescence spectroscopy with glycoconjugates

All fluorescence experiments were carried out as reported previously ^1, 24^ in 25 mM tris-HCl buffer at pH = 7.4. Thirty to forty μl (of 1 mg/ml) of protein solution was added to make-up the volume to 1 ml. This solution was used for the fluorescence studies. Emission spectra were recorded while exciting the protein solution at 280 nm. In a typical titration experiment, ligand solution of 2.5 μl (of 10 mM) was gradually added to the 1 ml of protein solution. Spectrum was recorded ranging from 290 nm to 520 nm after each addition. Buffer correction was done by subtracting the corresponding spectra, where, ligand was added to 1 ml of buffer solution containing no protein.

### Circular Dichroism (CD)

CD spectra were recorded at 25 °C on a Jasco-J-610 spectropolarimeter in 5 mM Tris-HCl buffer pH = 7.4 as reported previously ^1, 8, 25^. The spectra were recorded at a scan speed of 40 nm/min. All the measurements were made at concentration of 0.85 – 1 mg/ml of protein solution in the far UV region. Glycoconjugates ranging upto 84 to 417 μM were titrated with protein solution. The spectra were corrected by subtracting the spectrum of the buffer from them.

### Glycoconjugates and preparation of bulk solutions

Fixed amount of glycoconjugates was dissolved in 20 – 40 μl of dimethyl sulfoxide (DMSO). Buffer, 25 mM Tris buffer pH = 7.4, was added to make-up the volume to 1 ml giving the bulk solution of 10 mM. Glycoconjugate stability was monitored by UV-visible spectroscopy as reported earlier ^1, 8^.

### Protein ligand docking studies

All the docking studies were performed using the AUTODOCK 4.0 program ^26^. The initial structures of three lectins complex with sugar were downloaded from PDB database (1LES, 2PEL, 1SBF). The protein file preparation, *viz*., removing sugar, making monomer, etc., were performed with Accelrys viewrlite pro software [www.accelrys.com). The ligand preparations were carried out with Gaussview software. The initial input for glycoconjugates were taken from crystal structures of sugars ^27–29^, and glycoconjugates ^30–31^ published from our group. The basic sugars galactose (1), lactose (2) and glucose (4) were taken from crystal structures of lectins 1LES, 2PEL and 1SBF, respectively ^27–29^. The 1c and 4c are made from published crystal structures ^30–31^; 1B and 4B made from 1C and 4C, respectively. The glycconjugate 1a and 2a, were made from 1 and 2 by replacing the −OH at C1 position with −NH2 group; 1b and 4b are made from 1c and 4c respectively. The glycoconjugate 3 and 3C are made from 4 and 4C. The glycconjugate 2a, 2b and 2c were made by transferring the amine-, salicyl-iminoand naphthyl-iminomoiety from 1a, 1b and 1c, respectively. The synthetic glycoconjugates were independently geometry optimized using Gaussian 03 at B3LYP/6-31G level of theory, and were used in the present docking studies ^32^. Using ADT tools, the nonpolar hydrogens of the lectins were merged to their corresponding carbons and partial atomic charges were assigned (kollman). The nonpolar hydrogens of the ligands were merged and the rotatable bonds were assigned. The lectin structure was used as an input for the AUTOGRID program. AUTOGRID performed a pre-calculated atomic affinity grid maps for each atom type in the ligand plus an electrostatic map and a separate de-solvation map present in the substrate molecule. The dimensions of the active site box that was placed at the center of the lectin were set to 110 Å × 110 Å × 110 Å with a grid spacing of 0.375 Å. Docking calculations were carried out using the Lamarckian genetic algorithm (LGA) ^33^. Initially, we used a population of random individuals (population size: 150), a maximum number of 2500000 energy evaluations, a maximum number of generations of 27000, and a mutation rate of 0.02. One hundred independent docking runs were done for the glycoconjugate. The resulting positions were clustered according to a root-mean-square criterion of 0.5 Å.

## ASSOCIATED CONTENT

### Supporting Information

Supporting Information is available for this article.

## AUTHOR INFORMATION

## ACKNOWLEDGMENT

CPR acknowledges the financial support from DST (SERB), CSIR and DAE-BRNS, and the DST (SERB) for J.C. Bose National Fellowship and IIT Bombay for Institute Chair Professorship. AK and VKH acknowledge CSIR for their SRF.

---

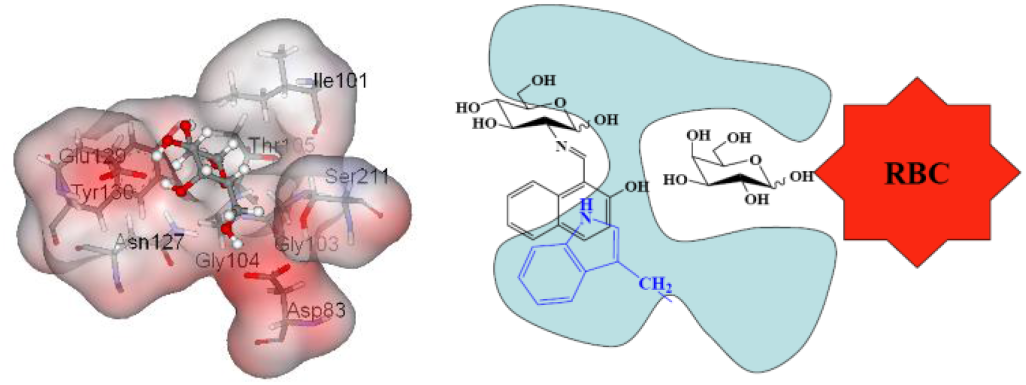

**SI 01**. Hemagglutination data

**Table S1:** Hemagglutination inhibition of lectin by glycoconjugates. Minimal inhibitory concentration shown for each lectins against glycoconjuagte.

| Conjugates | Lectins (Agglutination inhibition, mM) |  |  |  |  |  |  |  |
| --- | --- | --- | --- | --- | --- | --- | --- | --- |
|  | DLL-I | DLL-II | PNA | Moringa (MOL) | Unio | SBA | Lentil | Pea |
| Specificity→ | Glc/Man | Gal | Gal | Gal | Lac | Gal | Glc/Man | Glc/Man |
| **1** | NI | 3.9±0.01 | 7.8±0.03 | 3.9±0.02 | 7.8±0.02 | 31.25±0.7 | NI (500) | 125±5 |
| **4** | 62.5±4 | 125±5 | 500±10 | NI (500) | 15.6±3 | NI (500) | 62.5±5 | 31.25±1 |
| **3** | 15.6±3 | NI (500) | NI (500) | 250±10 | NI (500) | NI (500) | 15.6±3 | 15.6±3 |
| **2** | NI (500) | 15.6±3 | 0.97±0.03 | 15.6±3 | 0.48±0.01 | 62.5±5 | NI (500) | 62.5±5 |
| **1a** | NI (100) | 1.56±0.03 | 3.12±0.2 | 1.56±0.03 | 0.39±0.01 | 3.12±0.3 | NI (100) | 25±2 |
| **2a** | NI (100) | 12.5±0.3 | 0.39±0.01 | 1.56±0.03 | 0.048±0 | 3.12±0.3 | NI (100) | 50±5 |
| **2c** | NI (10) | NI (10) | 0.078±0 | 0.15±0.01 | 0.019±0 | 2.5±0.02 | NI (10) | 2.5±0.02 |
| **2b** | NI (10) | NI (10) | 0.15±0.01 | 1.25±0.02 | 0.039±0 | 2.5±0.01 | NI (10) | 5±0.01 |
| **1c** | 1.25±0.02 | 0.62±0.01 | 0.312±0 | 0.039±0 | 0.62±0.01 | 0.62±0.01 | NI (10) | 1.25±0.02 |
| **1b** | NI (10) | 1.25±0.02 | 1.25±0.02 | 0.312±0.01 | 2.5±0.02 | 2.5±0.02 | NI (10) | 5±0.01 |
| **4c** | 2.5±0.02 | NI (10) | NI (10) | NI (10) | NI (10) | NI (10) | 10±0.5 | 0.312±0 |
| **4b** | 5±0.01 | NI (10) | NI (10) | NI (10) | NI (10) | NI (10) | 20±0.5 | 5±0.01 |
| **4c** | 0.156±0 | NI (10) | NI (10) | NI (10) | NI (10) | NI (10) | 5±0.01 | 0.625±0.01 |
| ‘NI’ refers to no inhibition. |  |  |  |  |  |  |  |  |

SI 02: Fluorescence spectra obtained during the titration of the protein with the glycoconjugate.

**Figure S1.**
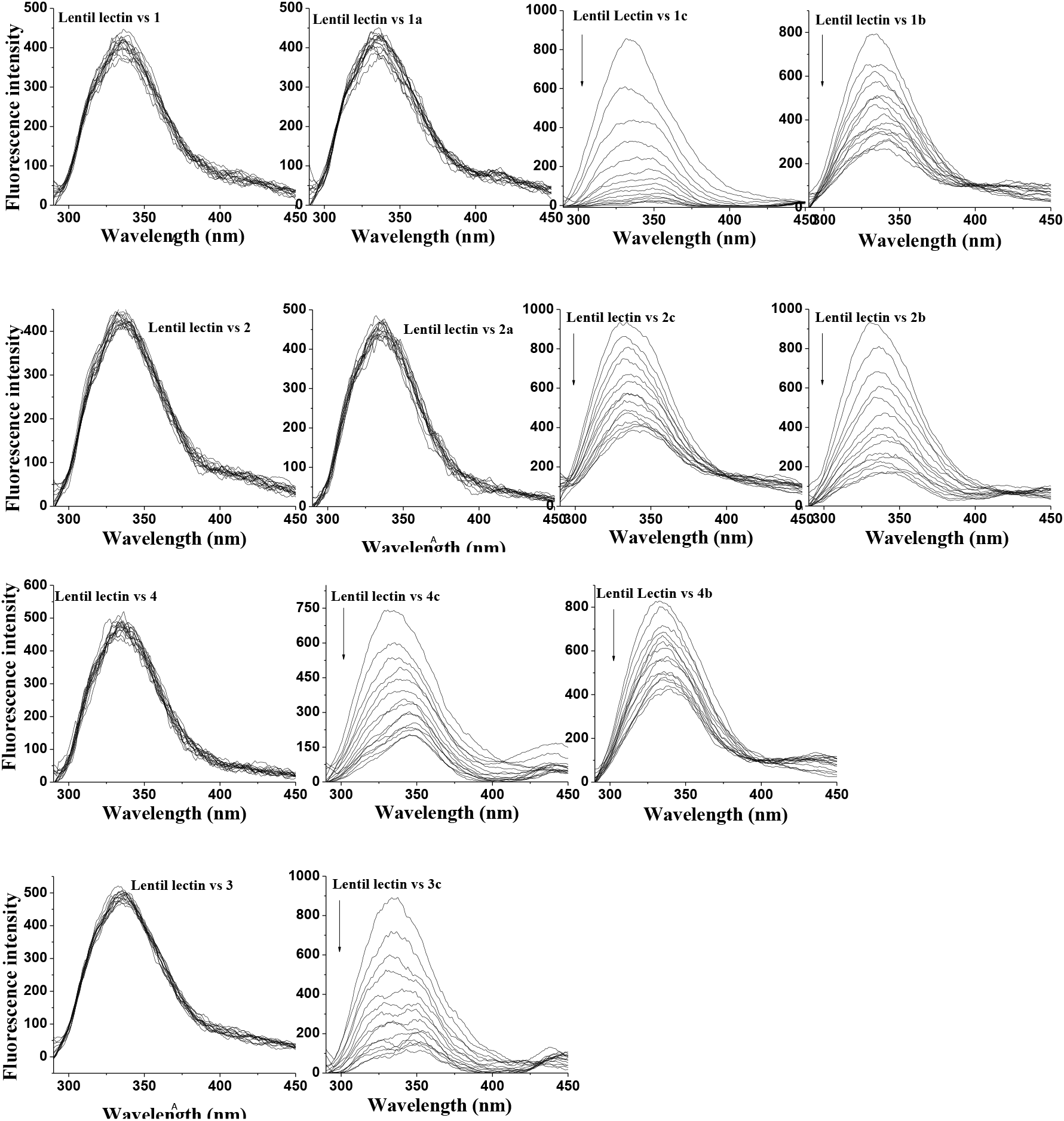
Fluorescence spectral traces obtained during the titration of lentil lectin with glycoconjugates.

**Figure S2.**
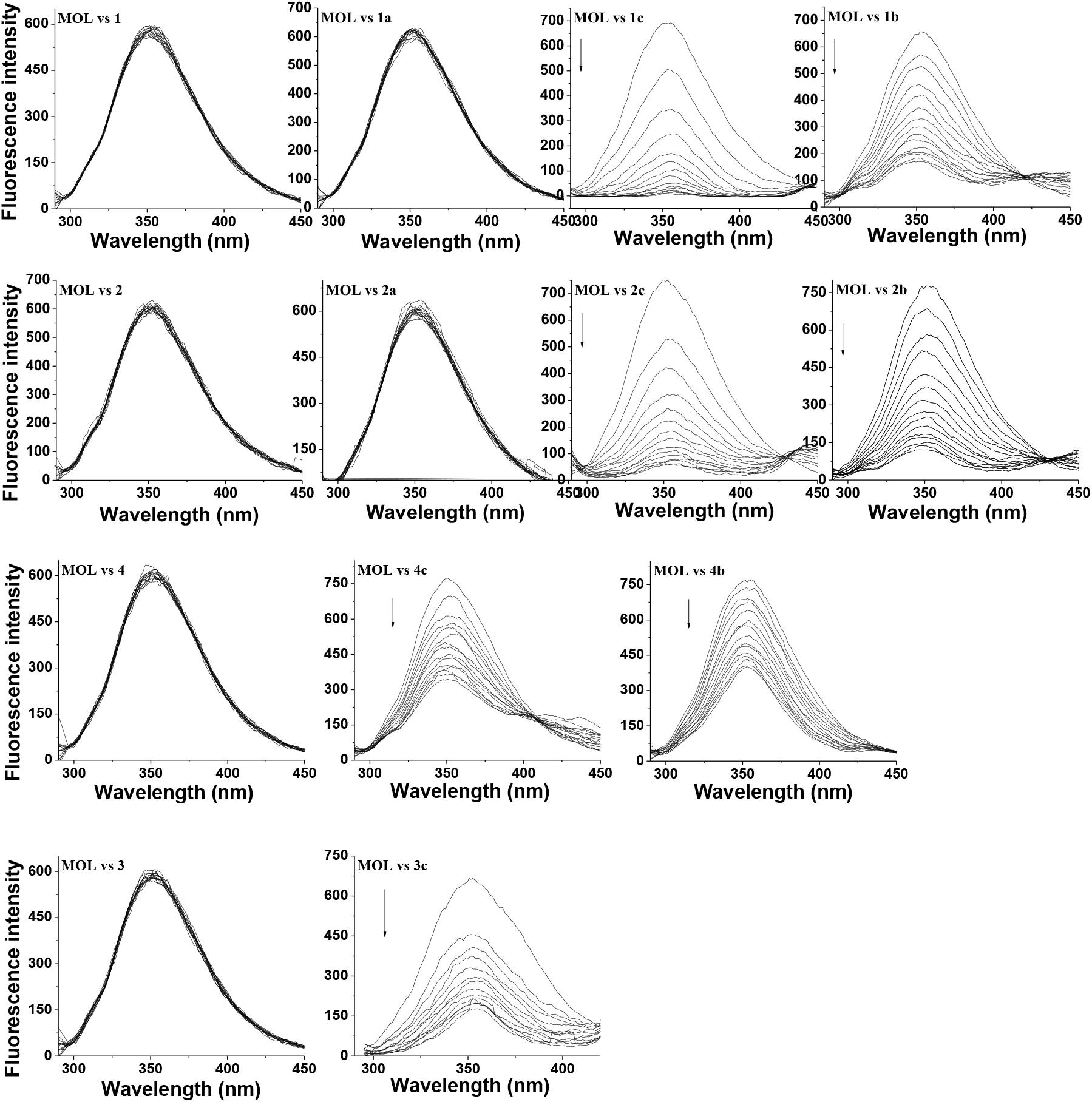
Fluorescence spectral traces obtained during the titration of MOL with glycoconjugates.

**Figure S3.**
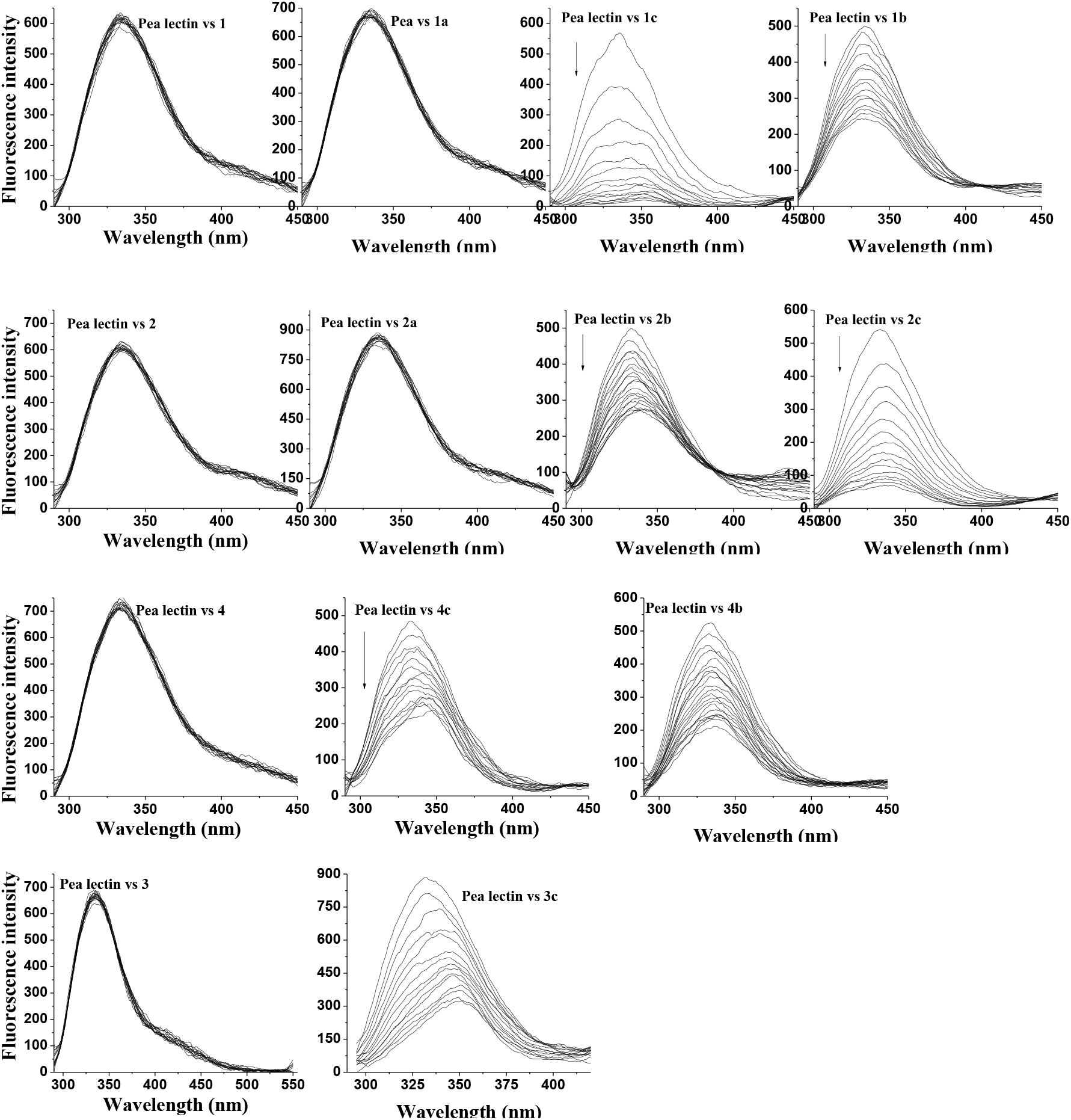
Fluorescence spectral traces obtained during the titration of pea lectin with glycoconjugates.

**Figure S4.**
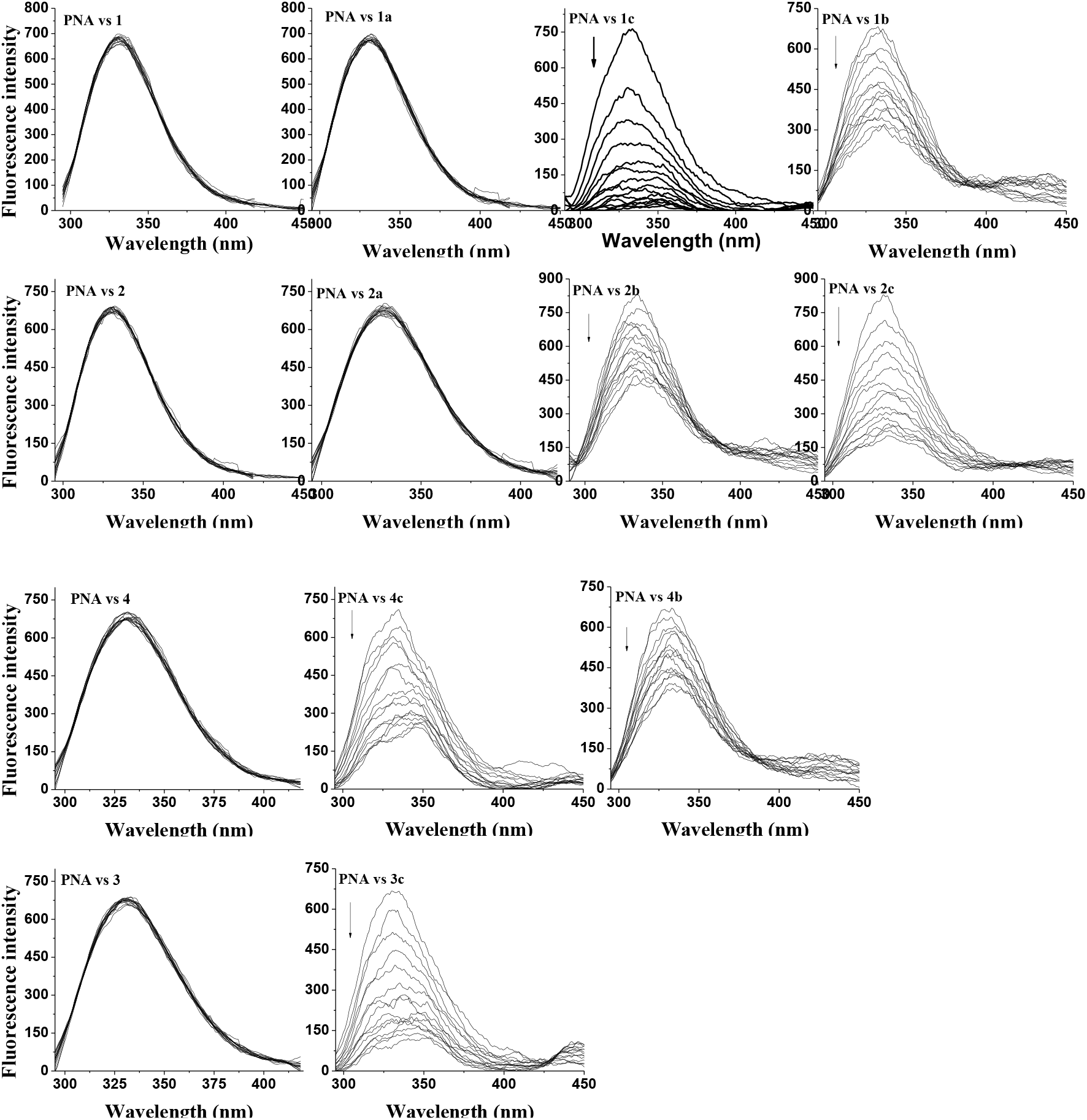
Fluorescence spectral traces obtained during the titration of PNA with glycoconjugates.

**Figure S5.**
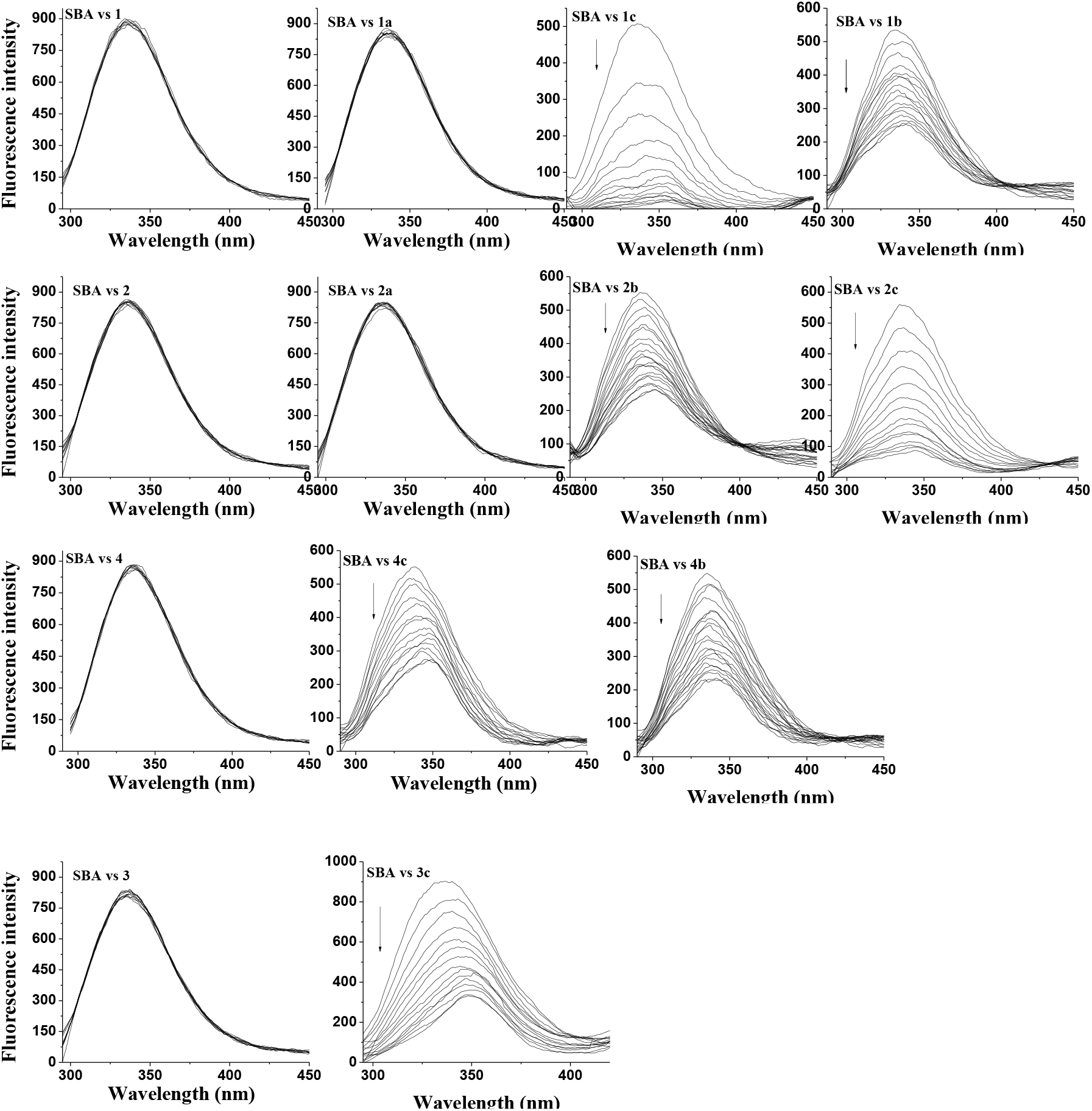
Fluorescence spectral traces obtained during the titration of SBA with glycoconjugates.

**Figure S6.**
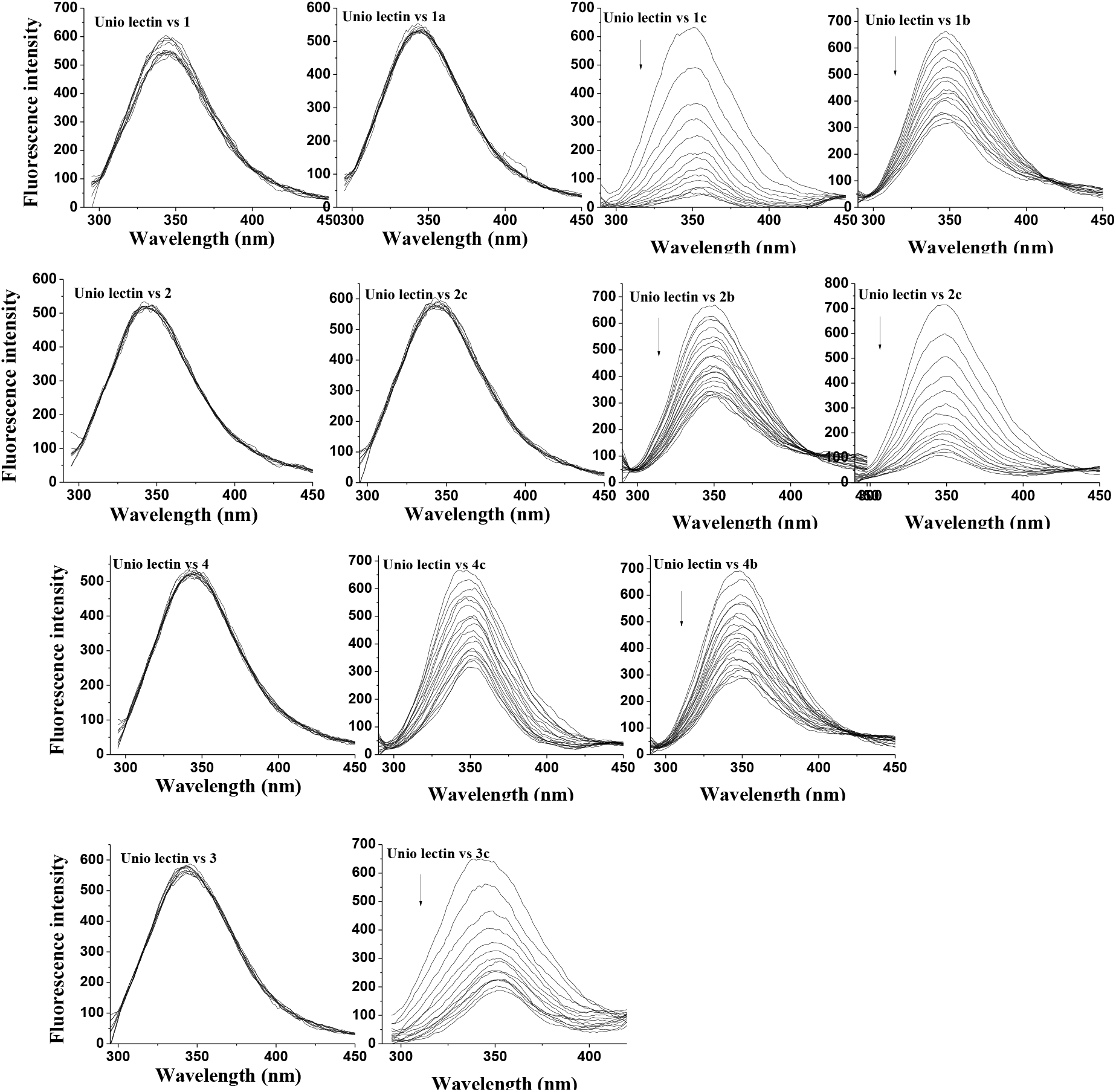
Fluorescence spectral traces obtained during the titration of unio lectin with glycoconjugates.

**Figure S7.**
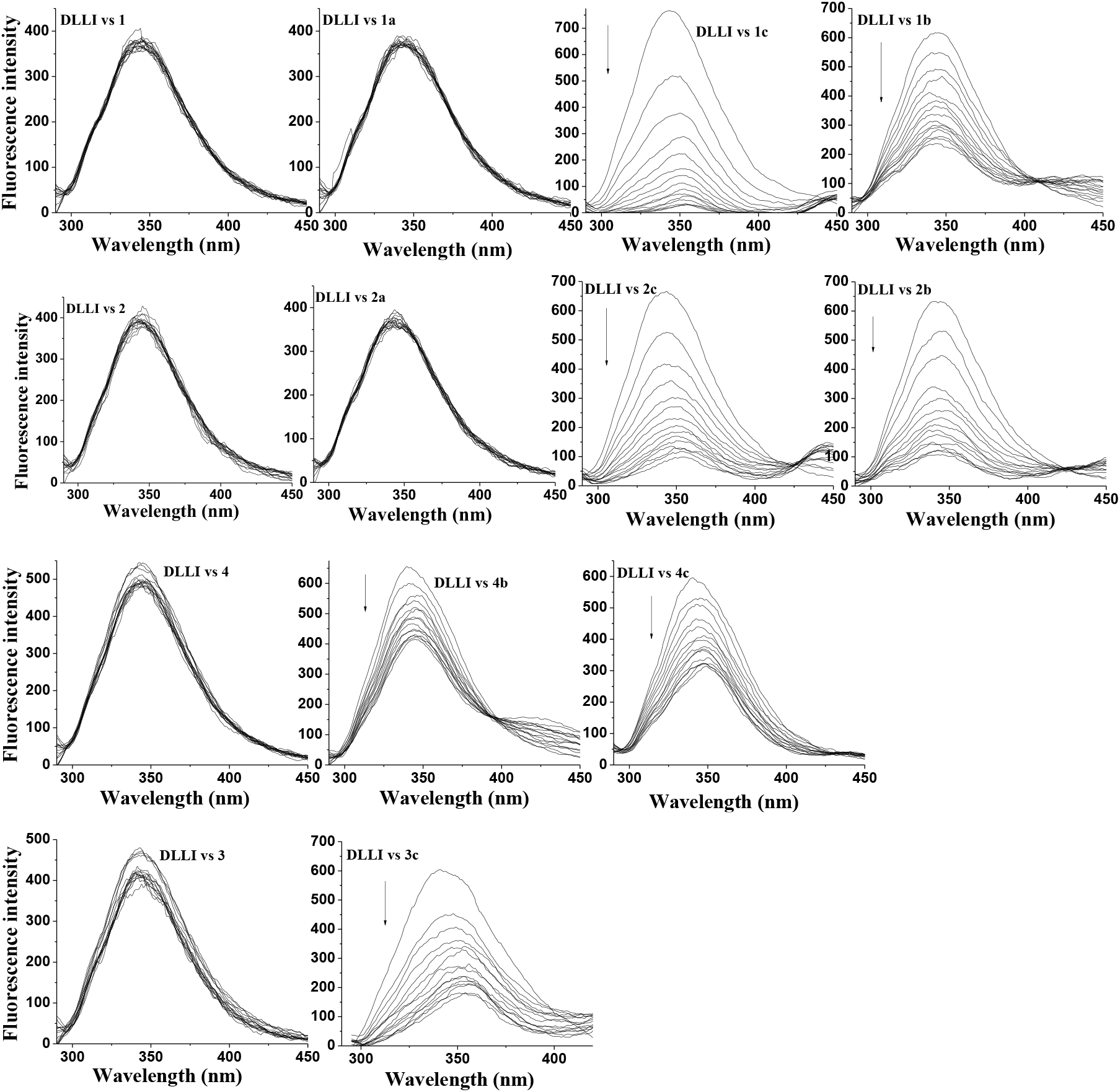
Fluorescence spectral traces obtained during the titration of DLL-I with glycoconjugates.

**Figure S8.**
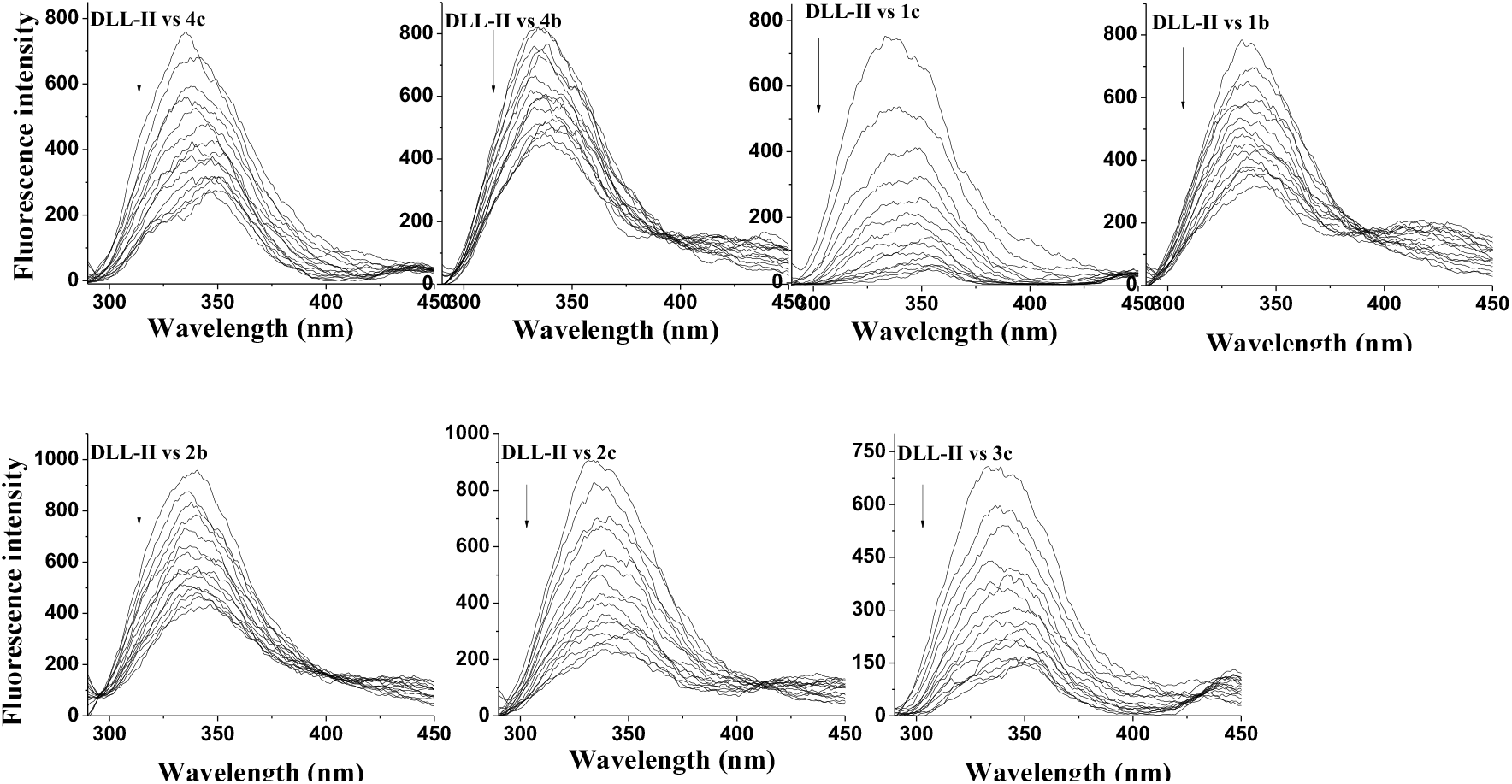
Fluorescence spectral traces obtained during the titration of DLL-II with glycoconjugates.

**Figure S9:**
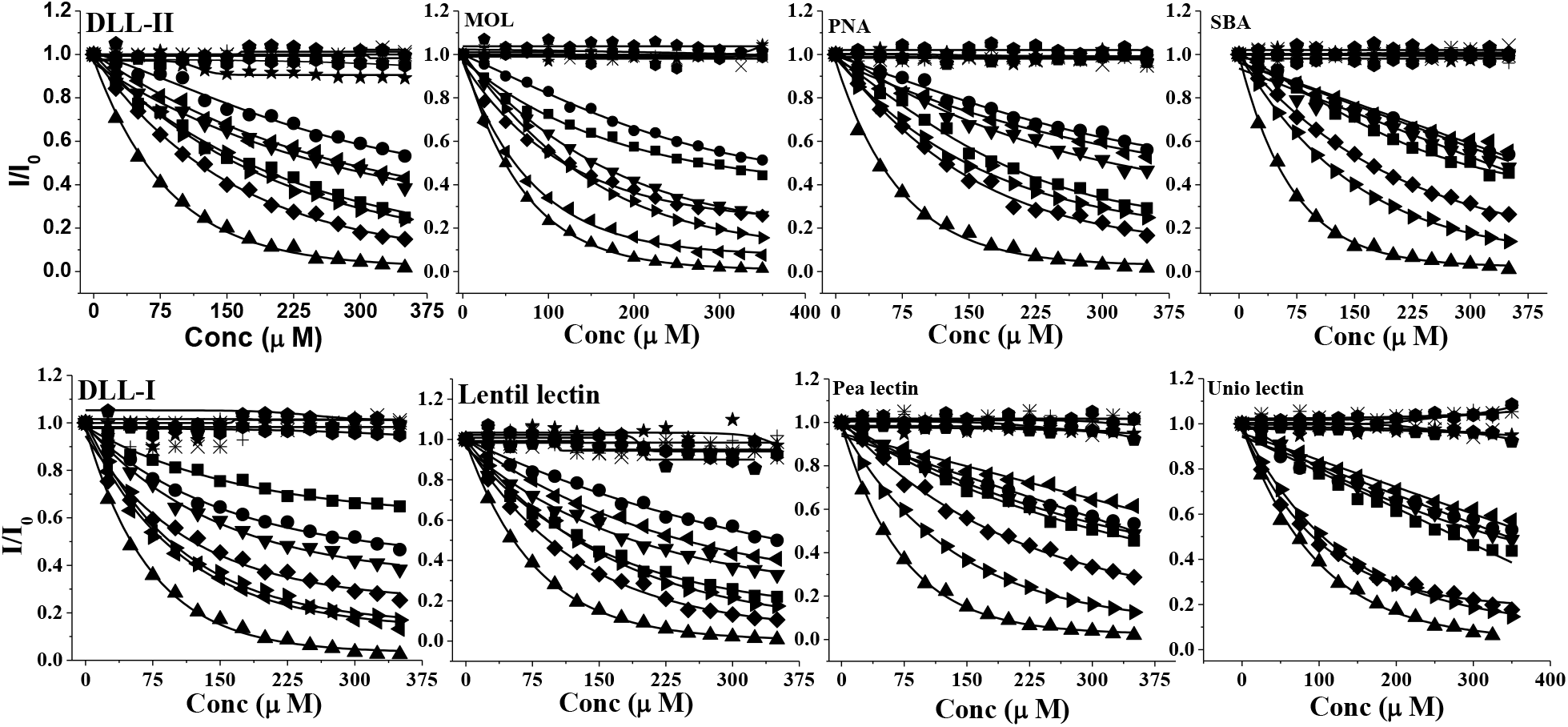
Relative intensity plots of different lectins, titrated with various glycoconjugates. (a) DLL-II, (b) MOL, (c) PNA, (d) SBA, (e) DLL-I, (f) Lentil lectin, (g) Pea lectin and (h) Unio lectin. Symbols have the same meaning in case of all lectins: ◆ – 1, – 1a, ▾ – 1b, ▲ – 1c, ✳ – 2, × – 2a, ▸ – 2b, ◂ – 2c, + – 3, ◆ – 3c, ★ – 4, ■ – 4b, ⚫ – 4c

**SI03**: Data, plots and trends based on the fluorescence studies.

**Table S2:** Trends of IF_50_ value, at the glycoconjugate point of view, as calculated from the relative fluorescence intensity plots. (Table foot notes same as in Table 1).

| Lectins | Trends in glycoconjugates IF<sub>50</sub> |
| --- | --- |
| DII-I | **1c > 2c = 2b > 3c > 1b > 4c > 4b** |
| Lentil | **1c > 3c > 2b = 4c > 1b > 2c > 4b** |
| Pea | **1c > 2b > 3c > 4c > 1b > 4b > 2c** |
| MOL | **1c > 2c > 3c > 2b > 1b > 4c > 4b** |
| SBA | **1c > 2b > 3c > 4c > 1b > 4b > 2c** |
| DLL-II | **1c > 3c > 2b = 4c > 1b = 2c > 4b** |
| PNA | **1c > 3c > 2b > 4c > 1b > 2c > 4b** |
| Unio | **1c > 3c > 2b > 4c > 1b > 4b > 2c** |

**Table S3:** Trends of IF_50_ value, at the lectins point of view, as calculated from the relative fluorescence intensity plots (Table foot notes same as in Table 1).

| Conjugates | Trends in lectins of IF<sub>50</sub> |
| --- | --- |
| **4c** | Lentil > DLL-II > PNA > DLL-I > MOL = Unio = SBA > Pea |
| **4b** | DLL-I > Lentil = MOL = Unio > Pea = SBA = DLL-II > PNA |
| **1c** | DLL-I = MOL = SBA = PNA > Lentil = Pea = DLL-II > Unio |
| **1b** | MOL > Lentil > DLL-I > DLL-II > PNA > Pea = SBA = Unio |
| **2c** | MOL > DLL-I > Lentil > DLL-II > PNA > SBA = PNA > Pea |
| **2b** | DLL-I > Pea = SBA = Unio > Lentil = MOL > PNA > DLL-II |
| **3c** | Lentil = Unio > MOL = DLL-II = PNA > DLL-I > Pea = SBA |

**Figure S10:**
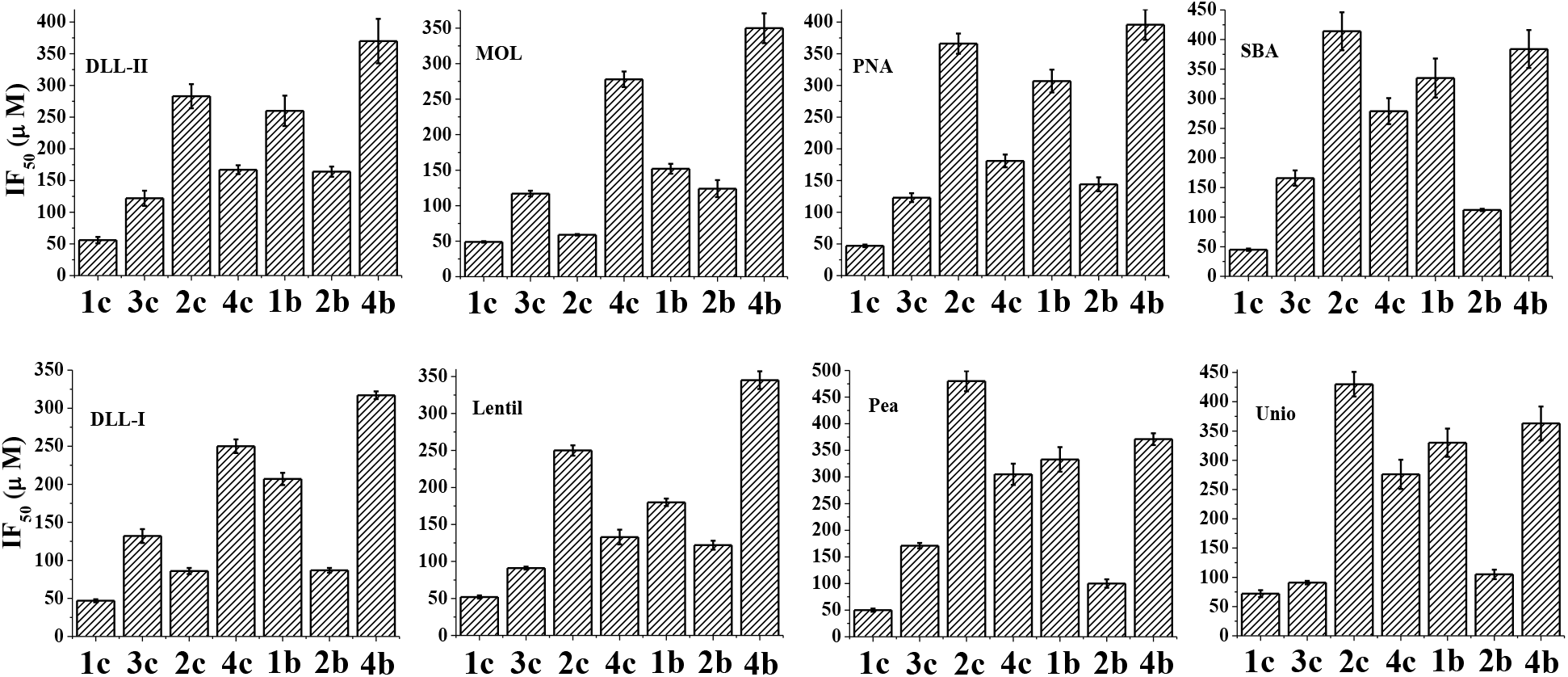
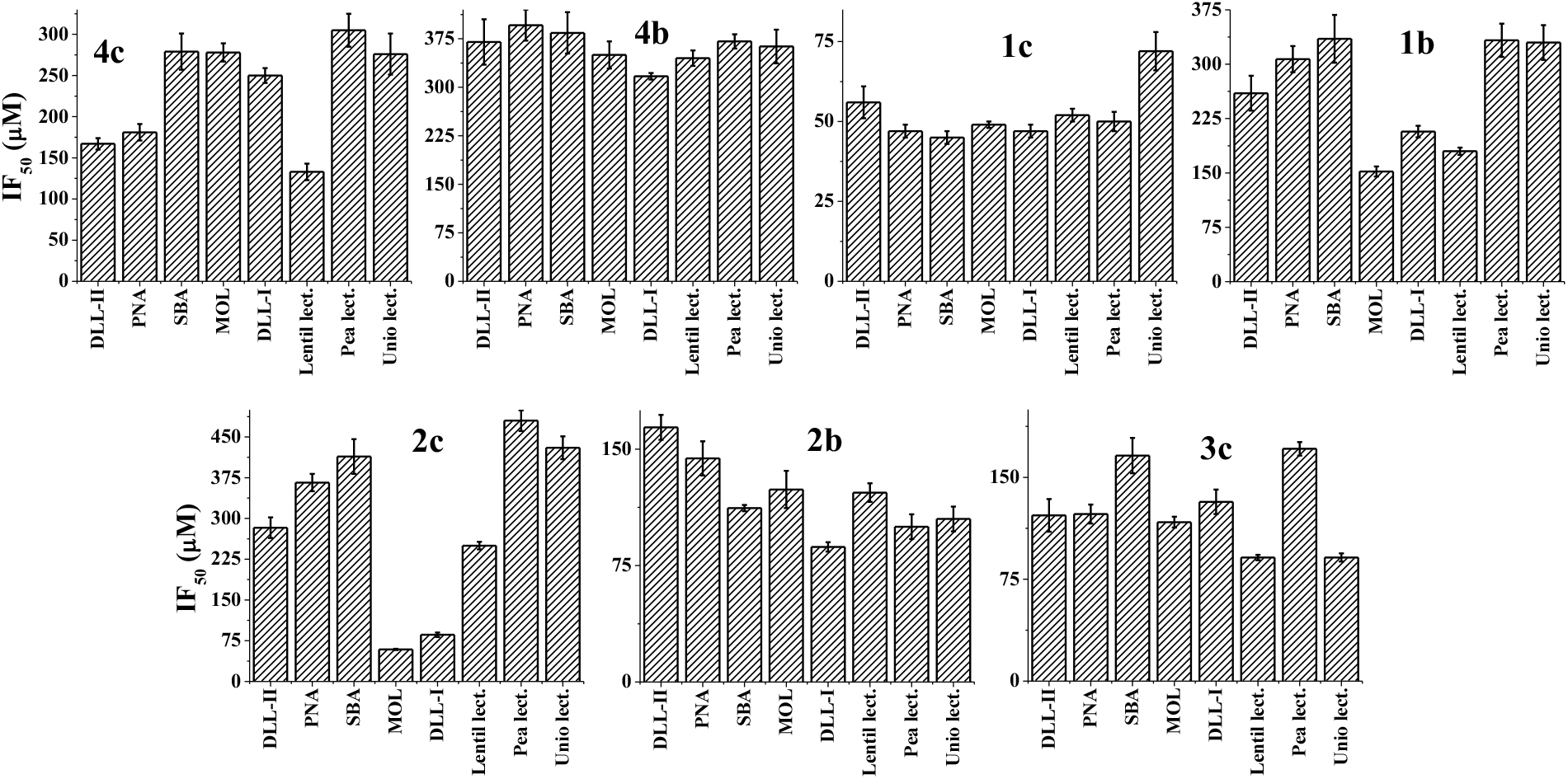
IF_50_ plots of lectins and glycoconjugate titration obtained based on fluorescence spectroscopy.

SI04: Stern-Volmer plots and the data.

**Figure S11:**
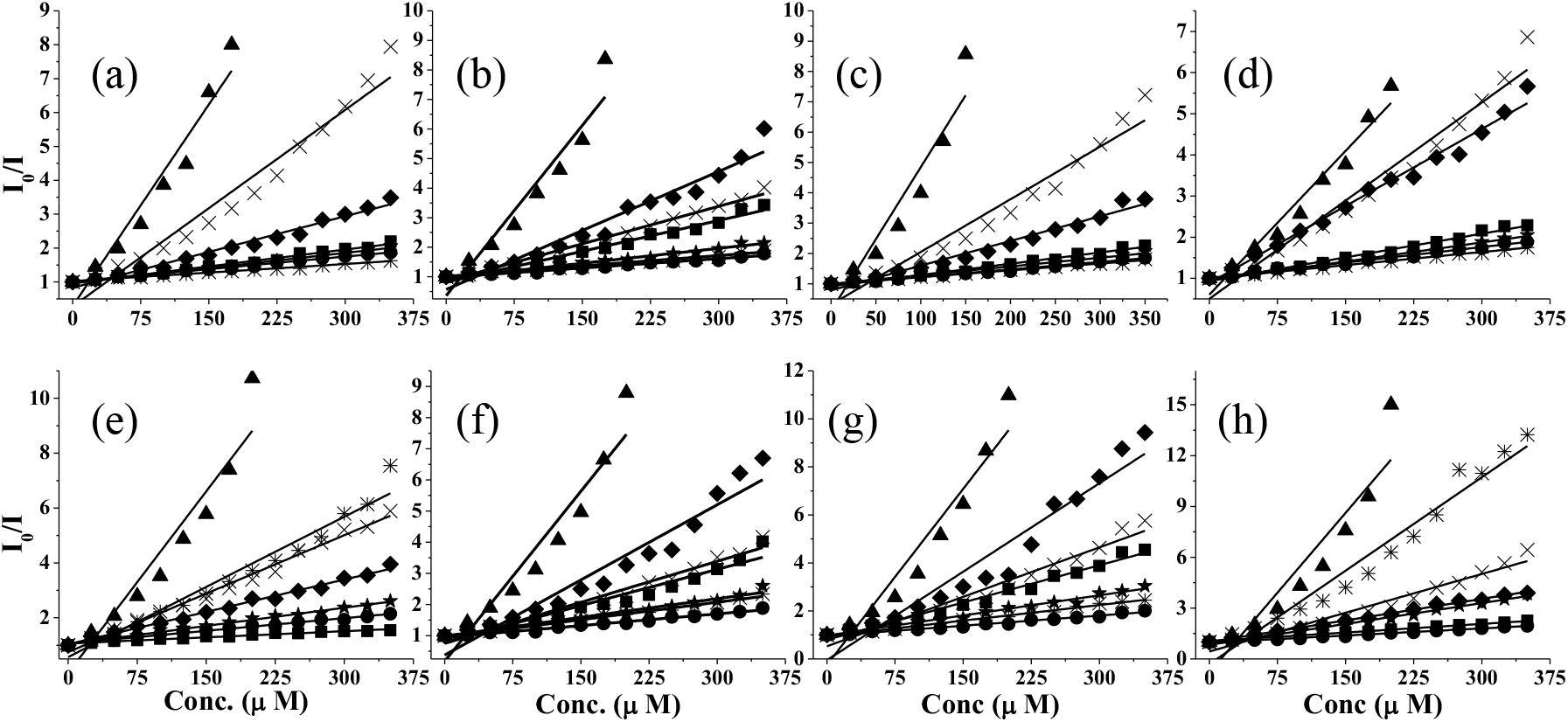
Stern-Volmer plots of different lectins, titrated with various glycoconjugates. (a) Pea lectin, (b) PNA, (c) SBA, (d) unio lectin (e) DLL-I, (f) DLL-II, (g) lentil lectin, (h) MOL. Symbols are same for all the plots: ★ – **1b**, ▲ – **1c**, × – **2b**, ✳ – **2c**, ◆ – **3c**, ⚫ – **4b** and ■ – **4c**.

**Table S4:** Different parameters calculated based upon the Stern-Volmer equation for different glycoconjugates which was titrated with the lectins. *K*_*sv*_ (M^−1^) – Stern-Volmet constant, *f*_*a*_ – fraction of fluorophore accessible to the quencher.

| Conjugates | $K_{sv} \times 10^3 (M^{-1})$ | | | | | | | |
| --- | --- | --- | --- | --- | --- | --- | --- | --- |
|  | DLL–I | Lentil | Pea | MOL | SBA | DLL–II | PNA | Unio |
| 4c | 1.4±0.01 | 10.3±0.3 | 3.3±0.1 | 3.0±0.1 | 3.7±0.01 | 7.7±0.4 | 6.9±0.1 | 4.3±0.2 |
| 4b | 3.1±0.01 | 2±0.1 | 2.7±0.1 | 2.0±0.1 | 2.8±0.1 | 2.5±0.1 | 2.1±0.1 | 2.98±0.1 |
| 1c | 36±2 | 36±0.3 | 41±0.04 | 49±3 | 37±2 | 26.3±1 | 31±0.1 | 21±0.2 |
| 1b | 4.4±0.2 | 5.5±0.2 | 2.9±0.1 | 8.0±0.2 | 3±0.1 | 4±0.1 | 3.3±0.1 | 3.1±0.1 |
| 2c | 17±0.9 | 4.3±0.1 | 1.7±0.3 | 36±2 | 2.5±0.6 | 3.7±0.1 | 2.3±0.1 | 2.3±0.1 |
| 2b | 14±0.5 | 13±0.7 | 19±0.01 | 15±0.2 | 17±0.1 | 8.7±0.4 | 8.4±0.3 | 16±0.1 |
| 3c | 18±3 | 24±2 | 12±0.3 | 18±0.4 | 16±1 | 15±0.1 | 13±0.8 | 14±0.5 |
| 1a, 2a, | ND | ND | ND | ND | ND | ND | ND | ND |
| 2, 1, 4, 3 | ND | ND | ND | ND | ND | ND | ND | ND |
| | $f_a$ | | | | | | | |
| **4c** | 0.89 | 0.88 | 0.87 | 0.79 | 0.88 | 0.86 | 0.82 | 0.83 |
| **4b** | 0.73 | 0.69 | 0.59 | 0.63 | 0.69 | 0.65 | 0.66 | 0.67 |
| **1c** | 0.92 | 0.87 | 0.88 | 0.79 | 0.89 | 0.88 | 0.79 | 0.88 |
| **1b** | 0.67 | 0.61 | 0.63 | 0.67 | 0.70 | 0.71 | 0.76 | 0.73 |
| **2c** | 0.88 | 0.83 | 0.79 | 0.77 | 0.81 | 0.89 | 0.91 | 0.93 |
| **2b** | 0.66 | 0.68 | 0.63 | 0.65 | 0.69 | 0.68 | 0.70 | 0.68 |
| **3c** | 0.88 | 0.91 | 0.89 | 0.81 | 0.88 | 0.87 | 0.80 | 0.84 |
| **1a, 2a,** | ND | ND | ND | ND | ND | ND | ND | ND |
| **2, 1, 4, 3** | ND | ND | ND | ND | ND | ND | ND | ND |

SI 05: CD spectra obtained during the titration of each of the lectin with all the glycoconjugates.

**Figure S12:**
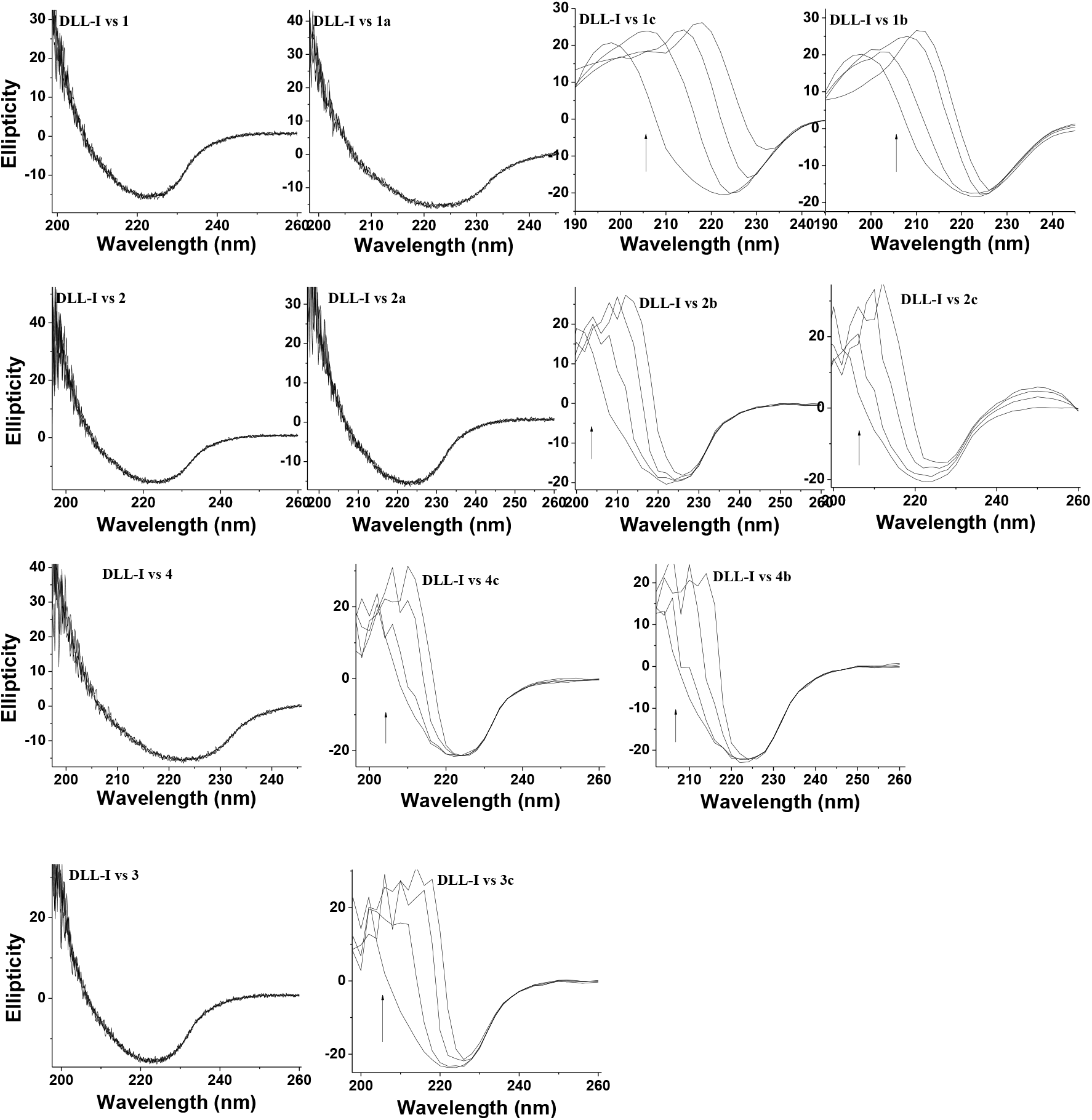
CD spectral traces obtained during the titration of DLL-I with glycoconjugates.

**Figure S13:**
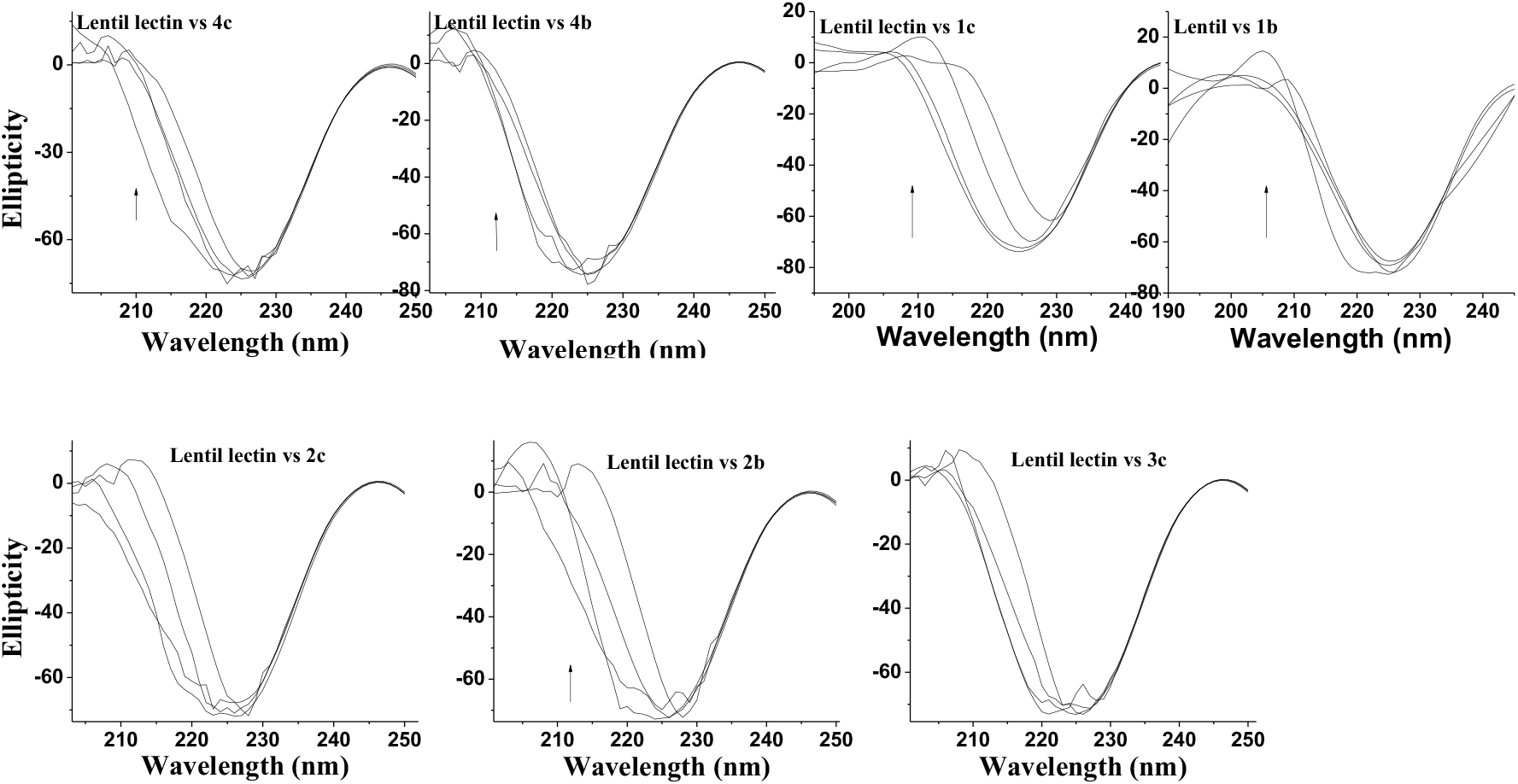
CD spectral traces obtained during the titration of lentil lectin with glycoconjugates.

**Figure S14:**
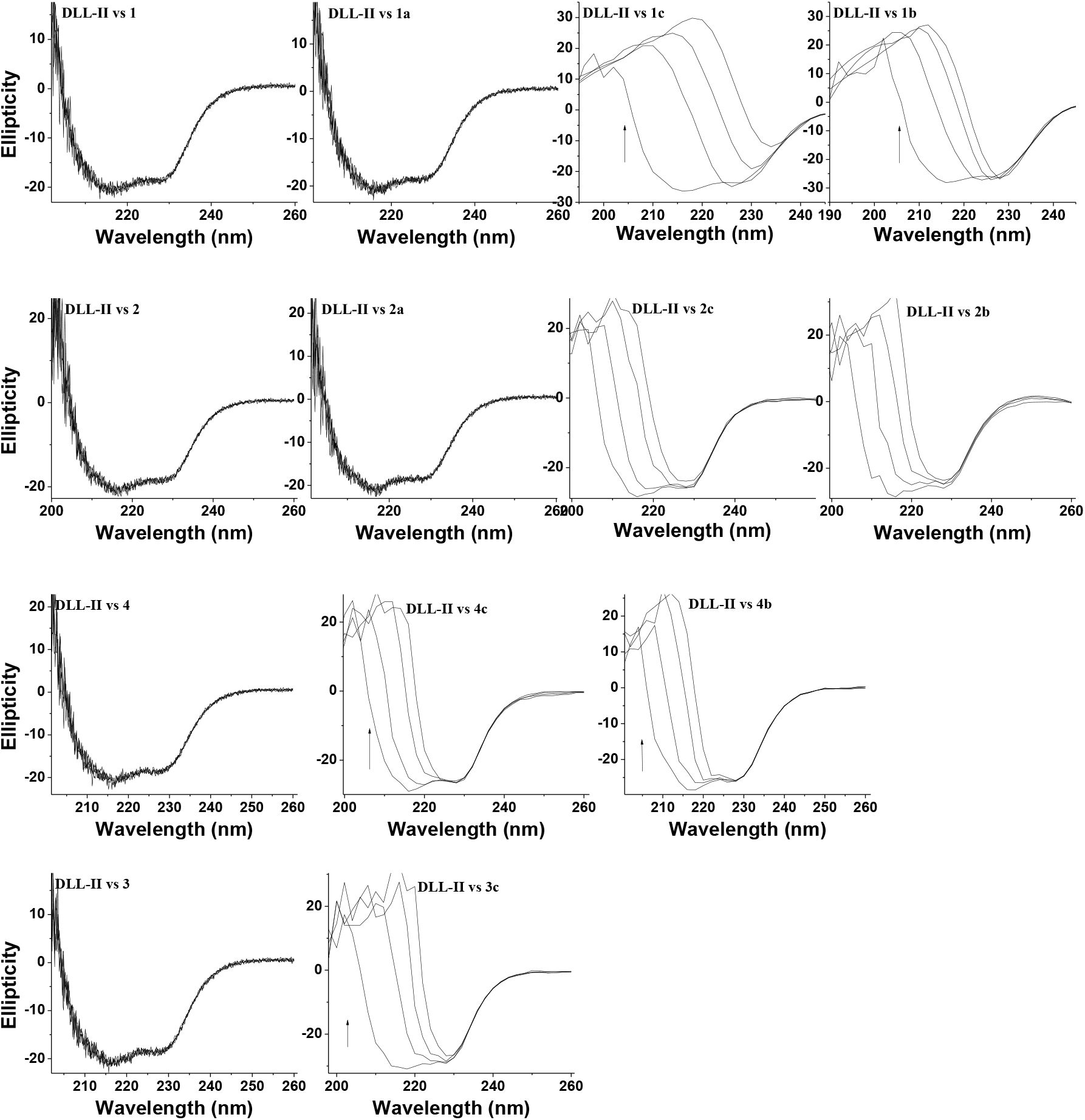
CD spectral traces obtained during the titration of DLL-II with glycoconjugates.

**Figure S15:**
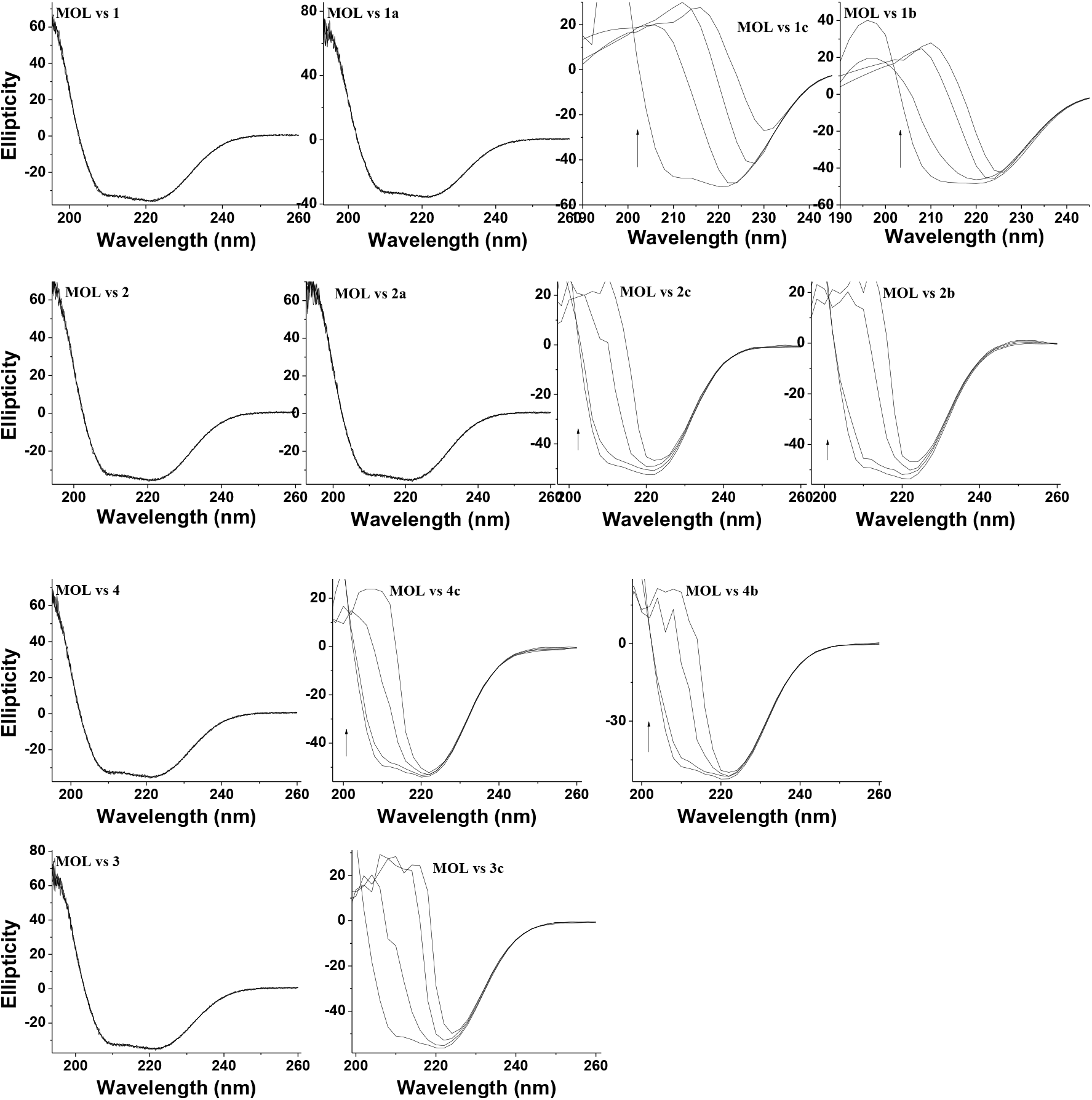
CD spectral traces obtained during the titration of MOL with glycoconjugates.

**Figure S16:**
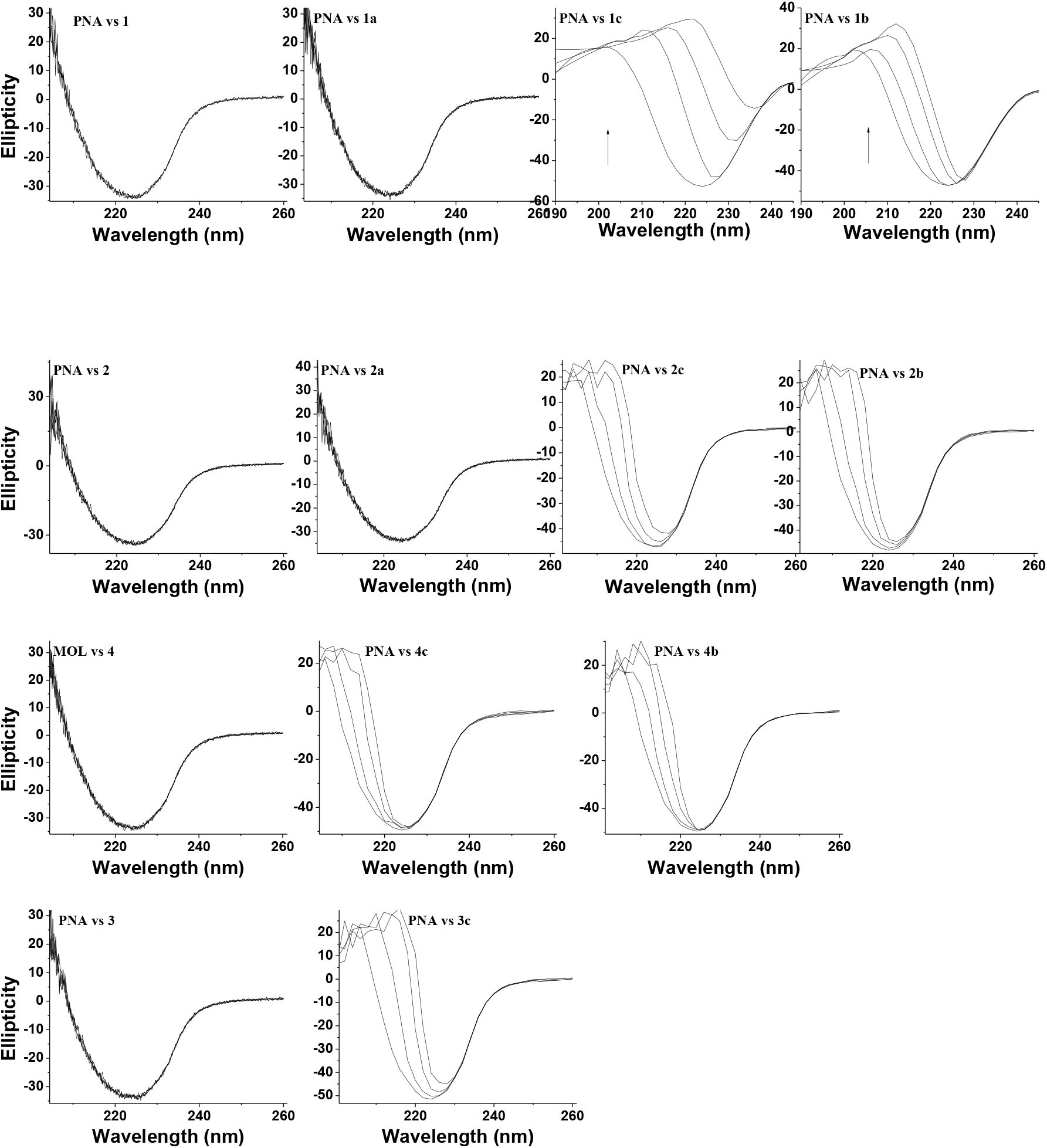
CD spectral traces obtained during the titration of PNA with glycoconjugates.

**Figure S17:**
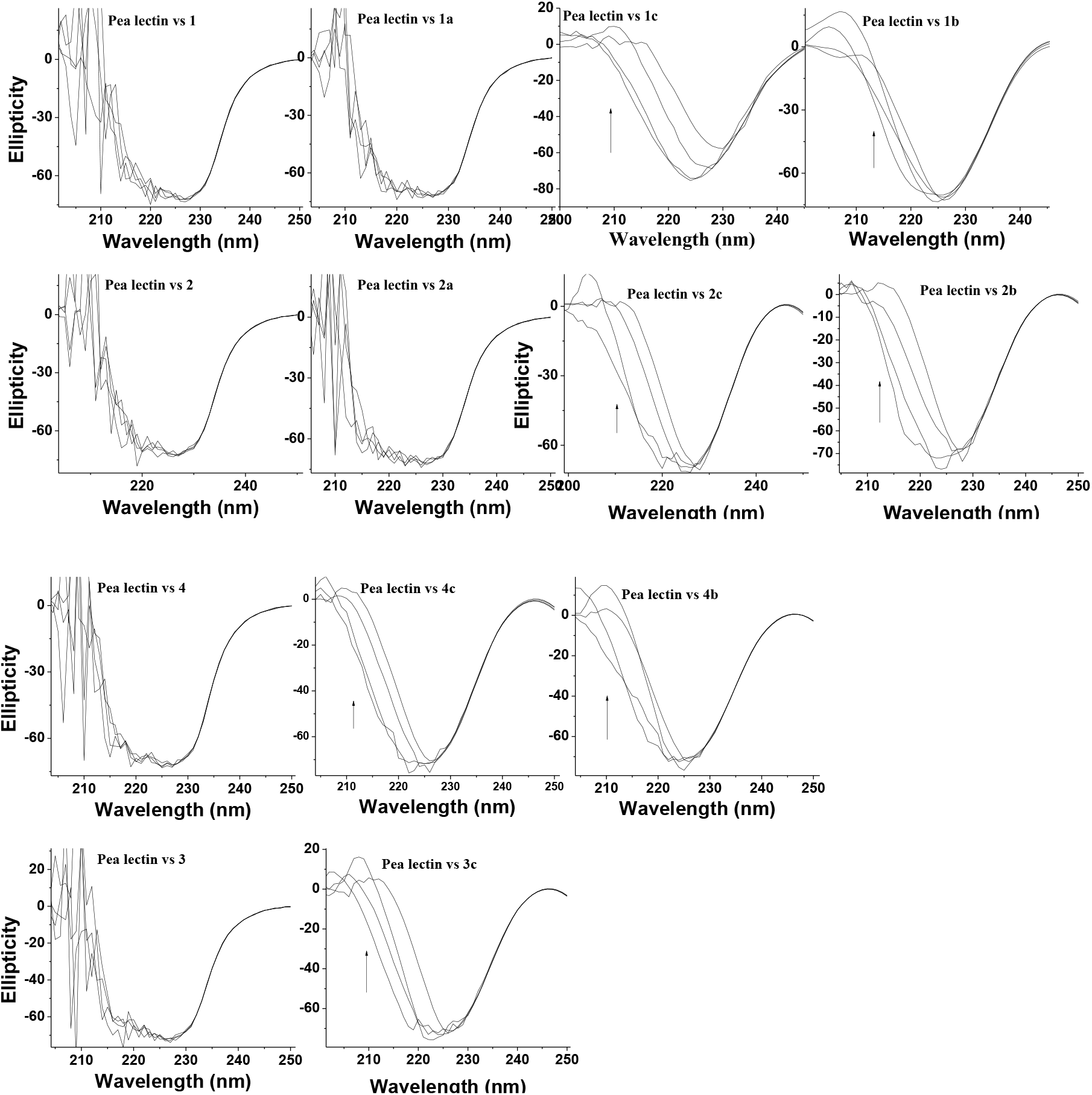
CD spectral traces obtained during the titration of pea lectin with glycoconjugates.

SI 06: Interaction data obtained from computational docking studies.

**Figure S18:**
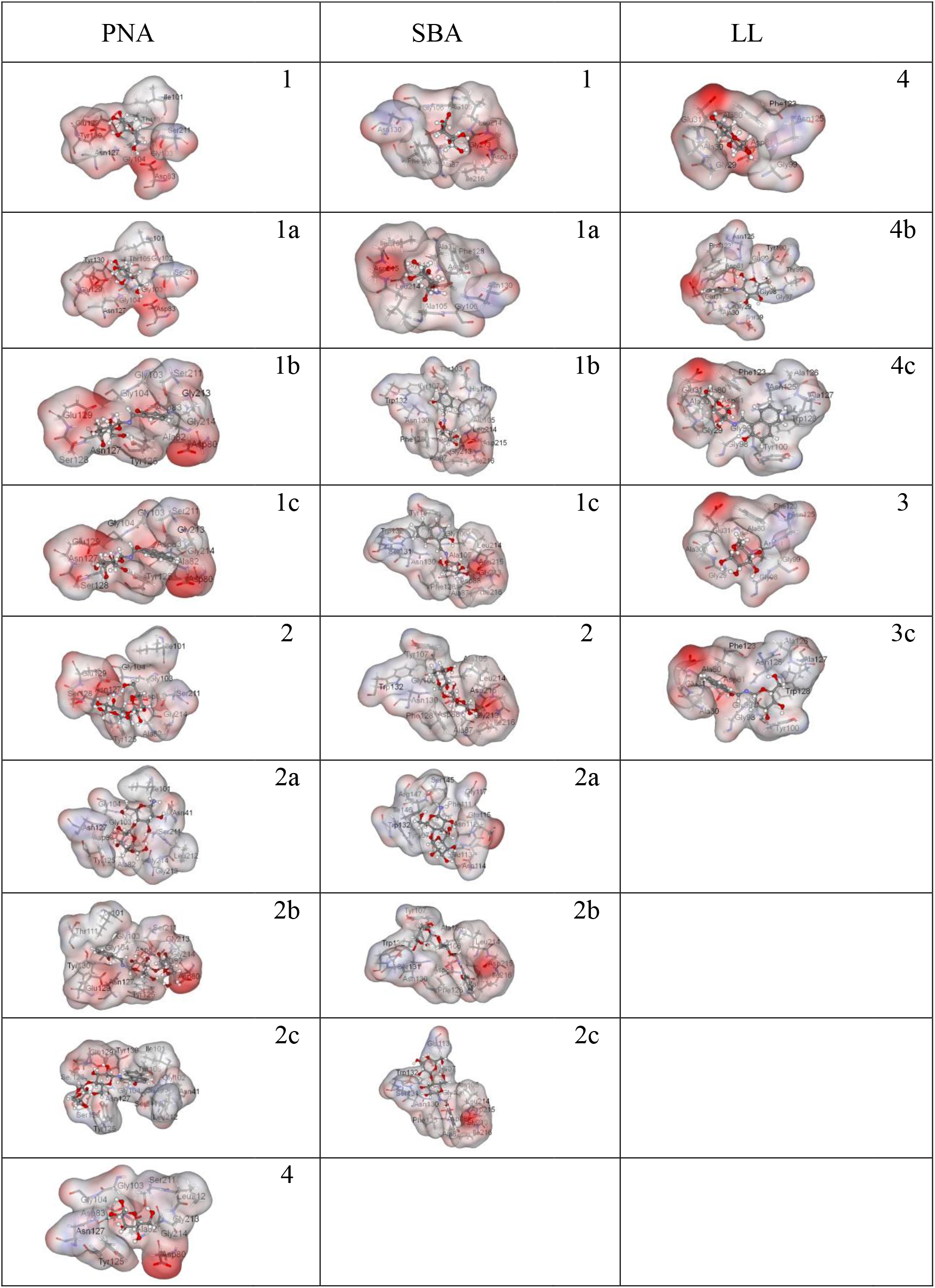
van der Waal surface representations (coloured with electrostatic potential) showing the binding of glycoconjugates at ***crd*** for PNA, SBA and lentil lectin.

**Figure S19:**
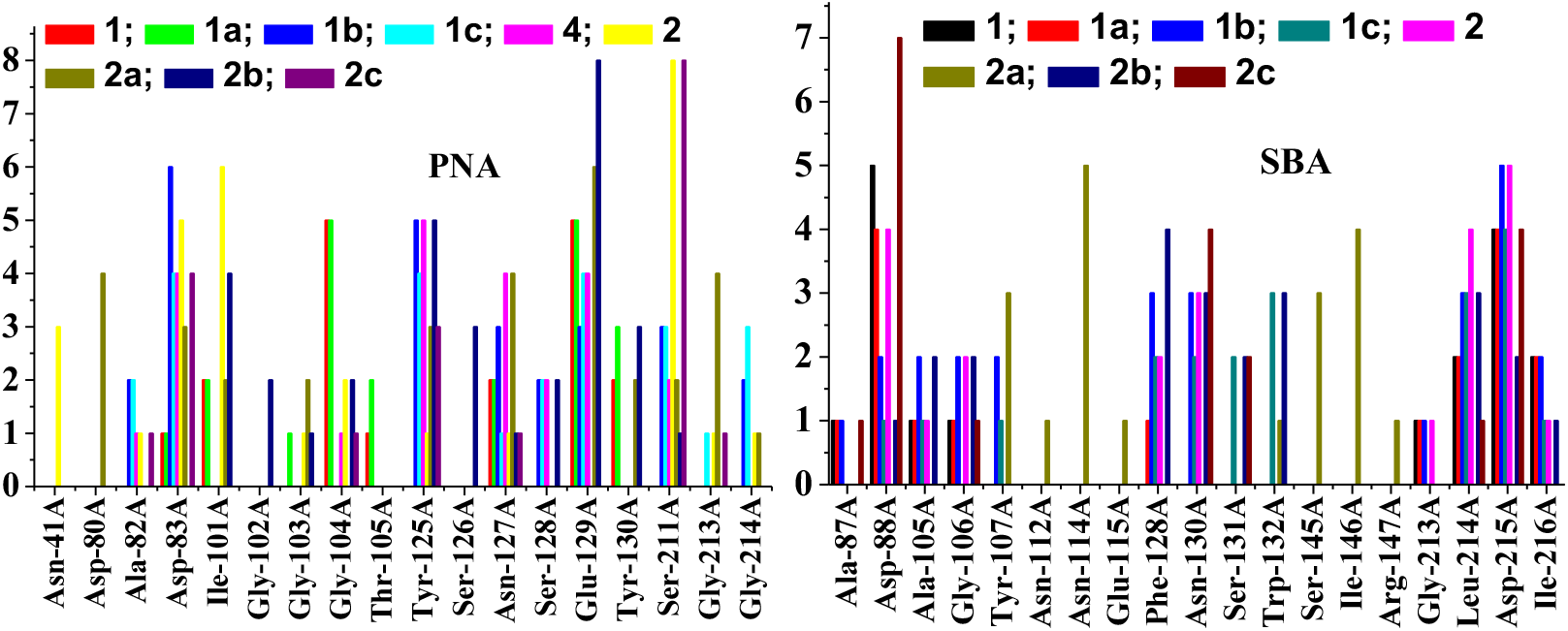
Residues specific interaction of lectin with glycoconjugates.

